# Full-length transcript architecture reveals RNA fate and regulatory state

**DOI:** 10.64898/2026.11.22.768006

**Authors:** Yifan Dai, Arnaud Stigliani, Jamie Yam Auxillos, William Garland, Kathryn Williams, Louis Souville, Benoît Ballester, Ane Amundarain, Mehdi Parviz, Anila George, Christian Vaagensø, Robin Andersson, Torben Heick Jensen, Albin Sandelin

## Abstract

Pervasive transcription generates diverse RNAs, including promoter-upstream transcripts (PROMPTs), coding and non-coding transcripts from protein-coding loci, and enhancer RNAs. How their full-length architectures relate to RNA fate remains poorly understood. Using ultra-deep sequencing of polyadenylated and non-polyadenylated capped RNAs from mouse embryonic stem cells, we show that nuclear exosome surveillance of PROMPTs is associated with transcript length and sequence composition. We show that for most protein-coding genes, capped transcripts are dominated by cytosolic 3′→5′ degradation intermediates or premature transcription termination products rather than full-length mRNAs. The relative abundance of intact mRNAs, cytosolic degradation intermediates and prematurely terminated transcripts varies between genes and is associated with gene architecture, expression level and biological function. Finally, we identify a class of convergently transcribed enhancers enriched for Polycomb repressive complex 2-associated proteins and poised enhancer features, with an increased propensity for activation during differentiation. Full-length transcript architecture thus reveals RNA fate and regulatory state.

## Introduction

Eukaryotic genomes are pervasively transcribed, producing a vast range of unstable long non-coding RNAs (lncRNAs), originating from promoters, enhancers and other regulatory elements^1–6^. Many of these transcripts are degraded by the exosome, a conserved 3′→5′ ribonuclease complex that functions in both the nucleus and the cytoplasm^7^. In the nucleus, substrate selection is mediated by adaptor complexes including the Nuclear EXosome Targeting (NEXT) complex and Poly(A) eXosome Targeting (PAXT) connection^8,9^. NEXT primarily targets non-polyadenylated RNAs, including promoter-upstream transcripts (PROMPTs)^8,10^, enhancer RNAs (eRNAs)^6^ and products of premature transcription termination (PTT)^11^, whereas PAXT acts on the polyadenylated isoforms of the aforementioned transcript classes and a subset of protein-coding mRNAs^8^.

Bidirectional transcription from nucleosome-depleted regions at promoters and enhancers generates PROMPT–mRNA and eRNA pairs, respectively^12^. At enhancers, eRNA transcription marks regulatory activity^6,13^. However, current models rely largely on short-read measurements of RNA fragments or ends^12^. Thus, relationships between full-length architecture, RNA surveillance, mRNA decay, PTT and enhancer state remain unclear. Long-read sequencing offers a route to address these questions^14,15^, but most approaches rely on poly(A) selection and therefore incompletely capture non-polyadenylated (pA^-^) RNAs that constitute a substantial fraction of nuclear transcripts. To this end, we previously developed TLDR-seq^16^, a long-read sequencing method capturing capped, full-length polyadenylated (pA^+^) and pA^-^RNAs.

Here, we combine TLDR-seq with nuclear RNA surveillance pathway depletions to characterize transcript architectures across promoters, protein-coding genes and enhancers in mouse embryonic stem cells (mESCs). We show that NEXT sensitivity is strongly associated with transcript length and sequence composition, and that NEXT-stabilized transcripts are frequently stochastically polyadenylated. We find that, for most mRNA genes, the majority of capped transcripts do not correspond to intact mRNAs but cytosolic 3′→5′ degradation intermediates or PTT products. Finally, we identify a class of convergently transcribed enhancers enriched for Polycomb repressive complex 2 (PRC2) proteins and chromatin features associated with poised regulatory elements. Our results demonstrate that full-length transcript structure encodes information about RNA fate and regulatory state.

## Results

### PROMPT stabilization leads to stochastic polyadenylation

Using mESC lines in which RBM7 or ZCCHC8 carried a dTAG degron^16^, we rapidly depleted either core NEXT component and generated triplicate TLDR-seq libraries alongside six controls (Ctrl; mean 68.1 x 10^6^ reads/library; Fig.1A, S1A–C). Overall, 83.7% of reads initiated within ±1 kb of mESC DNase-hypersensitive-site (DHS) midpoints^17^, consistent with earlier CAGE analyses^18^. We grouped reads by strand and overlapping 5′-end DHS to form DHS-anchored transcript clusters (Fig.1B), normalizing abundance to mitochondrial reads (Fig.S1C). Because PROMPTs are typically NEXT substrates^10^, we first analyzed 5,837 PROMPT-associated TLDR-seq clusters, defined as reads initiating from the same DHS as an annotated mRNA TSS but on the opposite strand. These RNAs were predominantly short, largely unspliced and on average strongly stabilized upon RBM7 or ZCCHC8 depletion (Fig.1C, S1D–H).

**Figure 1.**
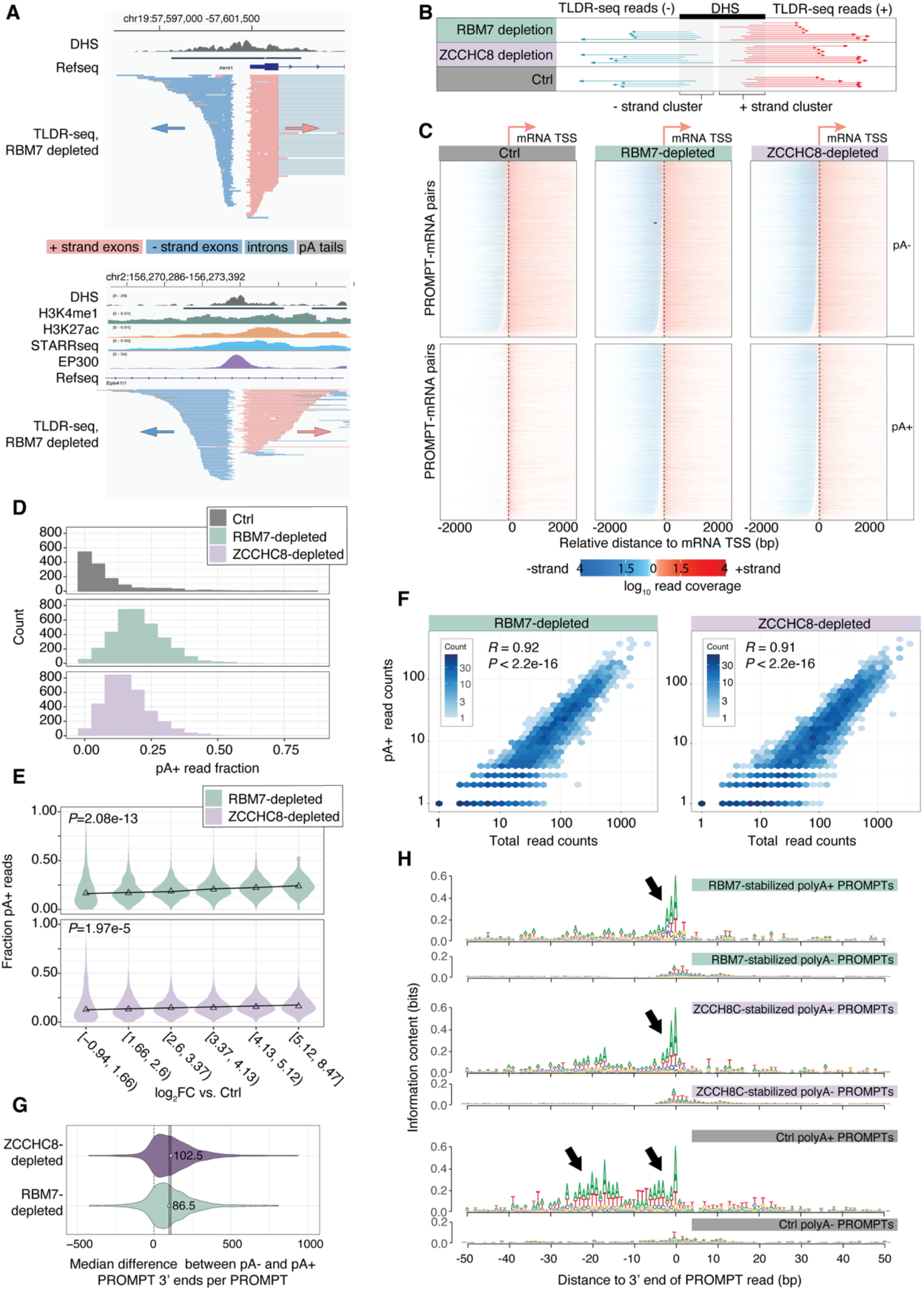
Stochastic polyadenylation of NEXT-stabilized PROMPTs. **A: Genome-browser examples of TLDR-seq data underlying the study.** Top shows a PROMPT-mRNA pair, bottom shows a putative enhancer region. The X axis shows genomic coordinates. mESC DNase-seq and (for the lower figure) STARR-seq and ChIP-seq read coverage tracks are shown, and RefSeq select gene models. TLDR-seq reads from RBM7-depleted cells are shown as horizontal bars, where blue indicate minus strand transcription, red indicates plus strand transcription (also shown by arrows) **B: DHS-anchored framework used for analysis.** TLDR-seq reads, shown as arrows where color indicates strand as in panel A, whose 5’ ends overlap DNase-hypersensitive sites (DHSs) are quantified on each strand for each treatment (rows). Each strand forms one TLDR-seq read cluster for further analysis, where reads in the cluster may have different 3’ ends, splicing and polyadenylation patterns. **C: PROMPT-mRNA pair heat map visualization.** Each row corresponds to one PROMPT-mRNA pair, oriented so that mRNAs are transcribed from left to right and PROMPTs right to left, as indicated on top. X axis shows distance to the mRNA TSS in bp. Color intensity shows TLDR-seq read depth, colored by strand. Reads are divided into two groups based on polyadenylation status. Data from Ctrl, RBM7- and ZCCHC8 depletions are shown. **D: Distributions of fraction polyadenylated (pA^+^) TLDR-seq reads from PROMPT clusters.** The X axis shows the fraction of pA^+^ reads in a TLDR-seq cluster. The Y axis shows the number of clusters. Color indicates TLDR-seq treatment: only clusters with more than five reads are shown. **E: Relation between PROMPT polyadenylation and stabilization following RBM7/ZCCHC8 depletion.** The Y axis shows the fraction of polyadenylated reads in TLDR-seq clusters assigned to PROMPTs. The X axis shows RBM7 or ZCCHC8 depletion log_2_FC vs. Ctrl of TLDR-seq clusters assigned to PROMPTs, divided into 6 bins. Colors indicate depletion type. Triangles indicate medians of distributions, connected by trend lines. ANOVA test *P* values are shown. **F: Relation between pA^+^ and total TLDR-seq read count for RBM7- or ZCCHC8-depletion-stabilized PROMPTs.** The Y and X axis shows number of TLDR-seq pA^+^ reads and total TLDR-seq reads, respectively in a given depletion (as indicated), in log_10_ scale. Hexagons are colored by the number of observations (log_10_ scaled). Pearson’s *R* and correlation test *P* values are shown. **G: Median read 3’ position difference of PROMPT pA^+^ and pA^-^ reads following NEXT depletion.** The X axis shows the median of the difference in genomic 3’ end positions between pA^+^ and pA^-^ TLDR-seq reads that are stabilized by RBM7 or ZCCHC8 depletions within each PROMPT. Positive X axis values indicate that pA^+^ reads are longer than pA^-^ reads, not counting the length of the pA tail, where 0 (dotted line) indicates no change. The Y axis shows the probability density distribution as violin plots. Lines, triangles and numbers within each distribution show distribution medians. **H: Sequence patterns around polyadenylated 3’ ends of PROMPT reads following RBM7 or ZCCHC8 depletions.** The Y axis in all sequence logos show information content in bits. The X axis shows the genomic distance in bp to the most used PROMPT 3’ end in each set. The top logo shows sequence patterns of dominant pA^+^ PROMPT 3’ ends stabilized by RBM7 depletion, while its paired logo just below shows the corresponding analysis on RBM7-stabilized pA^-^ PROMPT 3’ ends. Note that the Y axis scale on these plots are the same, but the lower is cut at 0.2 bits. The middle pair of logos show the same analysis on ZCCHC8-depletion-stabilized reads, while the last two logos show the same analysis for Ctrl library reads. Arrows indicate the canonical pA site (AWTAA) and the A/T enrichment patterns around 3’ ends

In agreement with a recent 3′-end based analysis showing post-transcriptional polyadenylation of stabilized NEXT substrates^19^, PROMPT RNAs stabilized after RBM7 or ZCCHC8 depletions were frequently polyadenylated (Fig.1C-D). The fraction of pA^+^ reads increased from 11% in Ctrl cells to 17% after ZCCHC8 depletion and 22% after RBM7 depletion, and the pA^+^ read fraction per PROMPT correlated with their degree of stabilization (Fig.1E). Restricting the analysis to PROMPTs lacking pA^+^ reads in Ctrl cells, pA^+^ and total read abundance were strongly correlated (Pearson’s *R*=0.91-0.92; Fig.1F), consistent with polyadenylation occurring as stabilized transcripts accumulate. However, pA^+^ reads extended further downstream: median pA^+^ 3′ end positions shifted by ∼86-107 nt relative to pA^-^3’ end positions (Fig.1G). Moreover, depletion-specific pA+ reads lacked the canonical upstream pA^+^ motif observed in Ctrl pA^+^ reads (Fig.1H, S1I). Instead, we observed A/T enrichment at the 3’ ends of pA^+^ reads compared to pA^-^3’ ends, suggesting polyadenylation was generated independently of canonical pA^+^ signals. Our results indicate that stabilization of PROMPT RNAs is frequently accompanied by stochastic post-transcriptional polyadenylation.

### PROMPT stabilization is correlated to RNA length and A/T content

We next asked which transcript features correlate with PROMPT sensitivity to NEXT-mediated decay. PROMPT stabilization after RBM7 or ZCCHC8 depletion increased strongly with transcript length (Fig.2A,B): <40% of short pA^-^ transcripts (0-100 nt) were stabilized, compared with >80% of 350-1,000-nt transcripts, with stabilization peaking at ∼600-700 nt. pA^+^ reads showed greater stabilization across all length classes, consistent with preferential polyadenylation of stabilized transcripts (Fig.1C-E).

**Figure 2.**
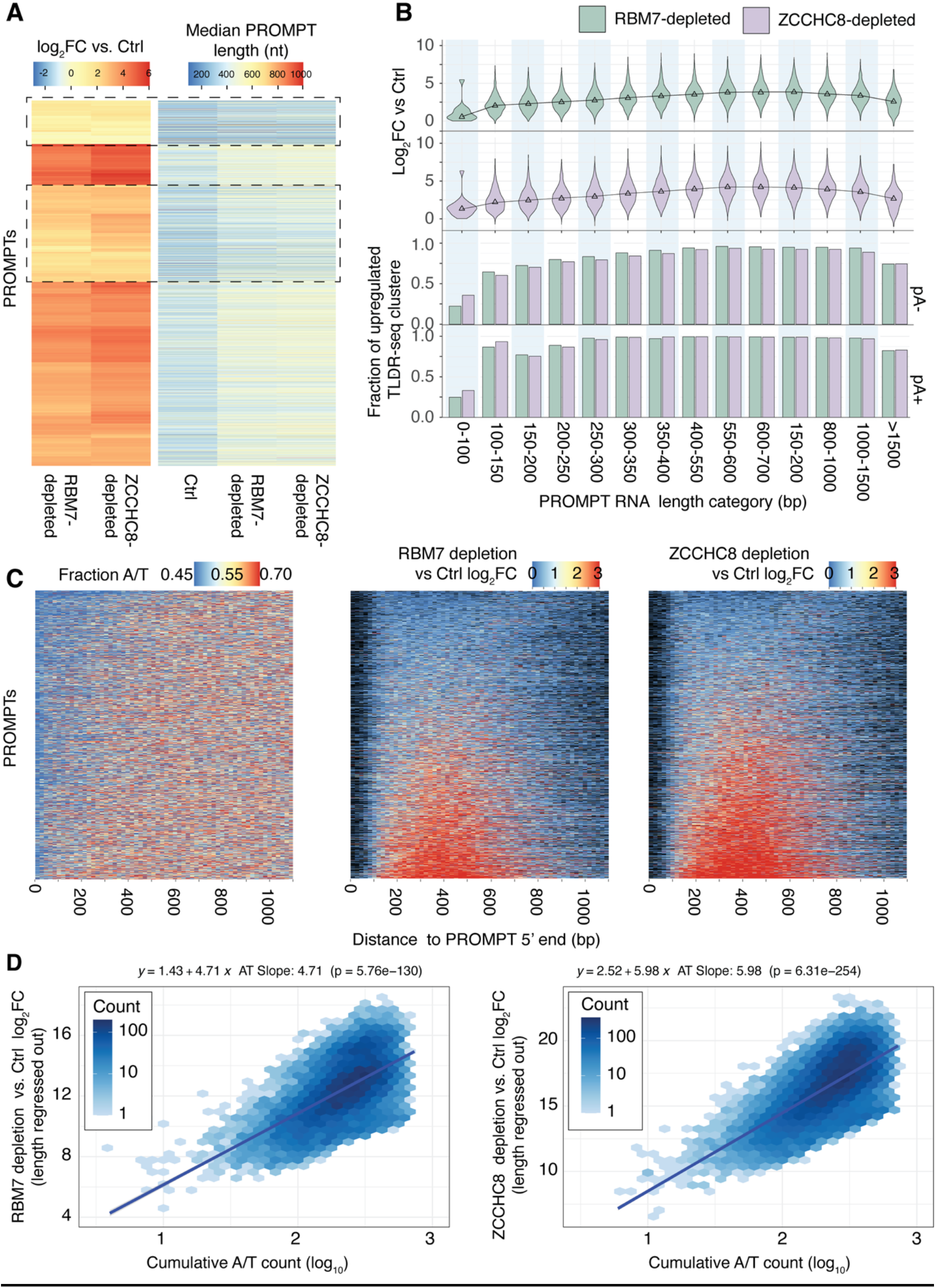
Association between PROMPT length, sequence composition and NEXT sensitivity. **A: Heatmap of PROMPT sensitivity to RBM7 and ZCCHC8 depletion alongside transcript length**. Rows represent TLDR-seq PROMPT clusters, hierarchically clustered based on RBM7 and ZCCHC8 log₂FC relative to Ctrl (first two columns). The remaining columns show median read lengths (nt) for Ctrl, RBM7, and ZCCHC8-depletion reads. Color intensity reflects log₂FC or median length values per TLDR-seq PROMPT cluster, as indicated on top. Dotted regions highlight clusters with the lowest log_2_FC values which typically have short median lengths (<300 bp). **B: Relationship between PROMPT length and sensitivity to RBM7 and ZCCHC8 depletions.** TLDR-seq reads within each PROMPT cluster were grouped into bins according to genomic length (x-axis). For each bin, log₂FC values (RBM7 depletion vs. Ctrl or ZCCHC8 depletion vs. Ctrl) was calculated for clusters with sufficient read coverage (Methods). The upper panel shows the distribution of log₂FC values per length bin (y-axis), with colors indicating depletion type. Median log₂FC values are shown as triangles and connected by a trend line. Lower panels quantify the number of clusters per bin exhibiting significant stabilization after depletion (log_2_FC >1 and *FDR*<0.05 vs Ctrl), stratified by pA⁺ and pA⁻ reads; bar colors reflect direction and magnitude of change. Only PROMPT clusters with >0 pA^+^ reads were analyzed. **C: Relationship between PROMPT A/T content and sensitivity to RBM7 and ZCCHC8 depletions.** Each row corresponds to one PROMPT TLDR-seq cluster. The X-axis represents consecutive 20-bp windows relative to the annotated PROMPT cluster 5′ end (leftmost bin: positions 0–19 bp, etc.). The left heatmap shows A/T fraction per bin. The middle and right heatmaps display RBM7 depletion vs. Ctrl and ZCCHC8 depletion vs. Ctrl log₂FC values, respectively, calculated using the reads whose 3′ ends fall within respective bin. Black bins indicate insufficient number of reads for robust log_2_FC calculation. Rows in all heat maps are hierarchically clustered by A/T content. **D: PROMPT sensitivity to RBM7 and ZCCHC8 depletions as a function of sequence composition independent of length.** Linear models were fitted using data from panel C to predict log₂FC (RBM7 or ZCCHC8 depletions vs. Ctrl) as a function of read length and cumulative nucleotide composition. The Y-axis shows length-adjusted log₂FC values (i.e., residuals after regressing out length effects). The x-axis represents cumulative A/T counts in the 20-bp genomic window in which the reads terminate (log₁₀ scale). Hexagons show the number of observations, and the blue line shows the fitted trend. The annotated ‘AT slope’ refers to the partial regression coefficient for cumulative A/T counts derived from the linear model, which quantifies the magnitude of the sequence effect when transcript length is held constant. Statistical significance of this sequence effect was assessed using a two-sided *t*-test.

Previous studies have identified A/T-rich pA^+^ motifs affecting exosome sensitivity^20–22^. Because promoter-proximal regions are G/C-rich whereas A/T content increases with distance from the mRNA TSS^4,23^, shorter PROMPTs are expected to be less A/T rich. Consistent with this, A/T content increased with distance from dominant PROMPT TSSs and positively correlated with RBM7 and ZCCHC8 stabilization, whereas extended A/T-poor regions exhibited little or no stabilization following depletion (Fig.2C). To determine whether this effect was statistically independent of transcript length, we modeled depletion stabilization as a linear function of RNA length and A/T content. For both RBM7 and ZCCHC8-depleted cells, sensitivity was strongly associated with A/T content after accounting for transcript length (slopes 4.51 and 5.21, respectively; *P* < 10⁻¹¹⁰; Fig.2D).

Together, these results show that NEXT sensitivity is independently associated with both transcript length and sequence composition. Short, promoter-proximal, G/C-rich PROMPTs were comparatively resistant to exosome targeting, whereas longer A/T-rich transcripts were preferentially degraded.

### Most protein-coding genes predominantly produce non-full-length transcripts

We next analyzed TLDR-seq reads initiating at DHSs overlapping annotated RefSeq promoters on the same strand, restricting the analysis to 5535 multi-exonic genes with one single promoter-overlapping DHS. Reads were classified by 3′ position as Full-length, First-exonic, First-intronic or Intermediate (ending elsewhere in the gene body; Fig.3A: see Methods); reads beyond gene boundaries or ending in 3′ UTRs distal to annotated 3’ends were excluded (1.9% of reads analyzed above extended across gene boundaries). Across genes, 22.2% were dominated (>50% of reads) by Full-length reads, 9.6% by First-exonic reads, 2.4% by First-intronic reads and 53.6% by Intermediate reads (Fig.3B; Figs.C-F show examples). Overall, 77.8% of genes contained more non-full-length than full-length reads, indicating that non-full-length RNAs constitute a large part of the steady state capped transcripts from most mRNA genes.

**Figure 3.**
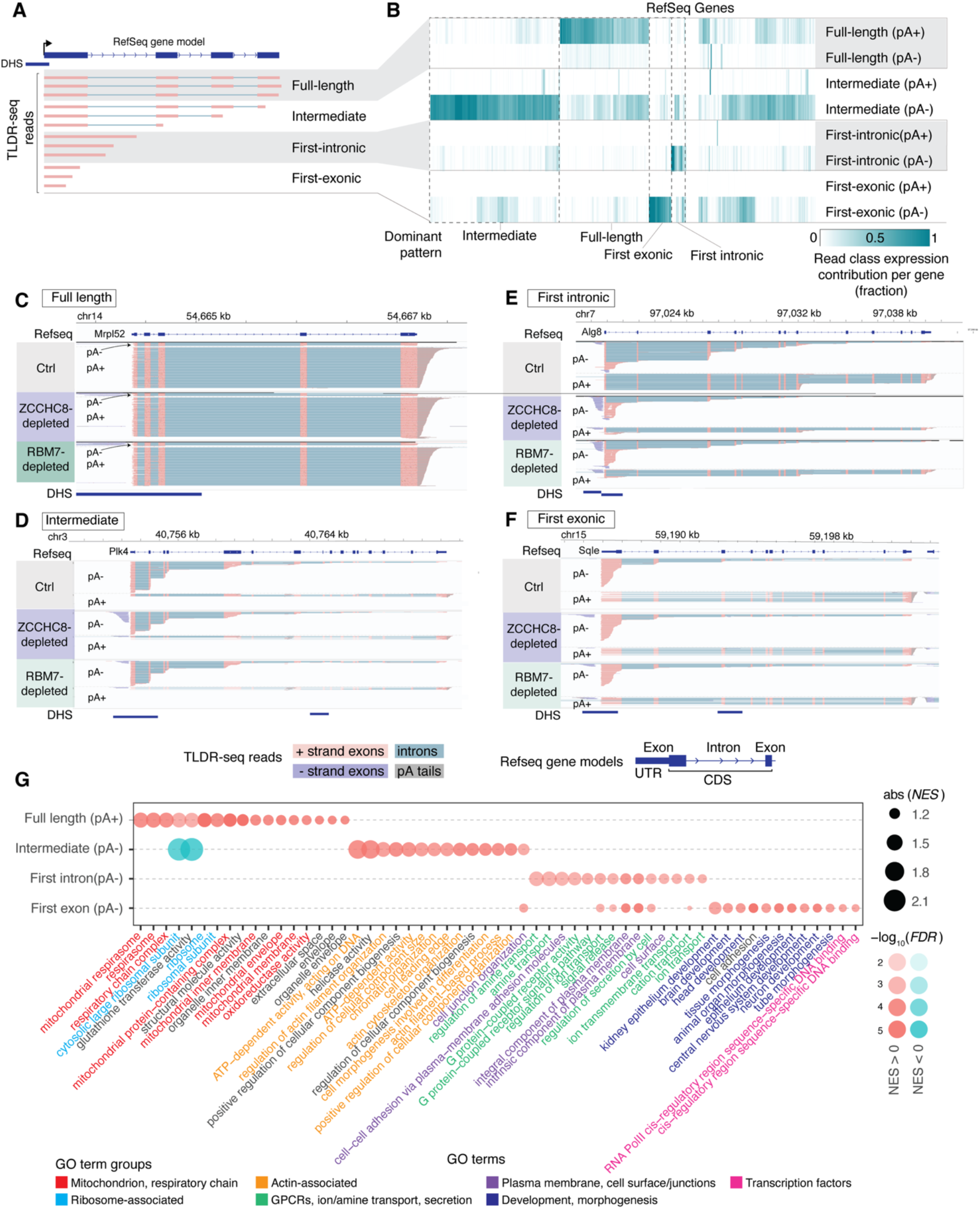
Analysis of genic TLDR-seq reads. **A: Classification scheme for genic TLDR-seq reads.** Schematic overview of the rules used to assign sense strand reads to classes for TLDR-seq clusters overlapping gene-associated DNase I hypersensitive sites (DHSs; blue). Classification is based on the genomic position of read 3′ ends relative to RefSeq gene models (top). Analysis was restricted to multi-exonic genes with a single transcribed DHS overlapping the annotated RefSeq promoter. Each read class (listed on the right) is illustrated with three representative TLDR-seq reads that meet the classification criteria. TLDR-seq exons are shown as red boxes and introns as blue lines. **B: Contribution of read classes to gene expression on a per-gene basis**. Columns indicate RefSeq genes. Rows correspond to read class as defined in panel A stratified by polyA status of reads. Heat map cell color indicates the fraction of total TLDR-seq gene expression that reads from each read class contribute with for a given gene (Ctrl cells, averaged across replicates). Columns are hierarchically clustered. The largest gene clusters dominated by one read class are highlighted. The small set of genes with 0 contribution from any class correspond to genes with high read contributions from excluded categories of reads (e.g. read-through). **C–F: Genome browser snapshots illustrating representative genes in which expression is dominated by one read class**. The X axis shows genomic coordinates (mm39), as indicated on top. RefSeq gene models are shown below: exons are blue boxes (larger boxes for coding sequence, smaller for UTRs), introns are shown as lines with chevrons indicating direction of transcription. TLDR-seq reads are shown from Ctrl, RBM7- and ZCCHC8-depleted cells: each row is one read, where plus strand exons are in red, minus strand exons are in deep blue, introns in light blue. pA^-^ and pA^+^ reads are separated as indicated on the left: for pA^+^ reads, the length of the pA tail is indicated in beige. mESC DHS peaks are shown in the bottom track **G: Gene set enrichment analysis (GSEA) relating gene ontology term enrichment with degree of expression contribution per gene from a given read class.** A GSEA analysis was conducted, assessing gene ontology (GO) term enrichment/depletion as a function of the fraction of reads per genes that each read class contributes with (data from panel B). Rows indicate read class as defined in panel A. Columns indicate GO terms. Dot size indicates absolute normalized enrichment score (NES), color indicates over/under-representation, transparency indicates associated *FDR* value. GO term descriptions are colored by overall functional supergroups.

To determine whether read-class composition was linked to gene function, we performed gene set enrichment analysis based on relative read-class contribution (Fig.3G). Genes enriched for Full-length reads were associated with energy production and protein synthesis, consistent with efficient production of mature mRNAs from broadly expressed housekeeping genes. In contrast, Intermediate-read enrichment was associated with cytoskeletal organization and cellular component biogenesis. First-exonic read enrichment was strongly associated with developmental regulation and transcription factors (TFs), whereas First-intronic read enrichment was associated with membrane, signaling, secretion and transport pathways. These associations indicate that read-class composition captures biological differences between gene groups, rather than only technical or abundance-related variation. This raised the possibility that class-specific transcript patterns are shaped by systematic differences in RNA biogenesis and gene architecture of functionally related genes, explored in the next section.

### Most non-full-length, genic transcripts arise from cytosolic 3′→5′ mRNA decay or early premature transcription termination

To investigate the origins of the transcripts from the read classes defined above, we generated TLDR-seq libraries from nuclear and cytosolic fractions of wild-type mES cells (Fig.S2A), and total RNA from ZFC3H1- and hMTR4(SKIV2L2)-depleted cells, to assess sensitivity to PAXT degradation (ZFC3H1) and joint NEXT-PAXT degradation (hMTR4; Fig.S2B,C). Because differences in RNA composition complicate direct normalization between compartments, we calculated nuclear to cytosolic expression ratios for reads belonging to each read class within each gene and compared the resulting ratio-rank distributions across genes (Fig.4A).

**Figure 4.**
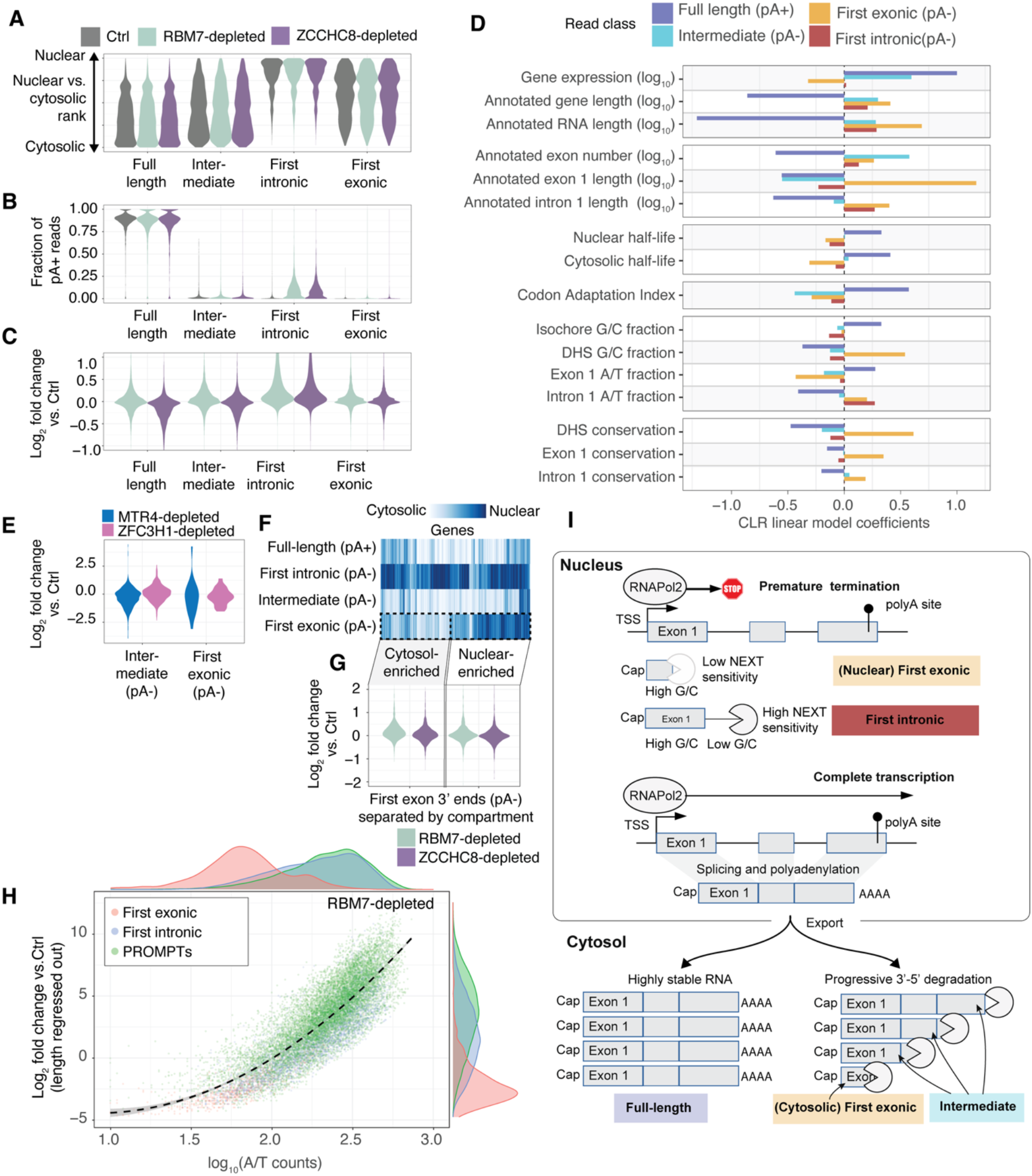
Properties of genic TLDR-seq read classes. **A: Nuclear-cytosolic ratio enrichment per read class.** Violin plots show the distribution of nuclear-cytosolic ratio (Y axis) for each read class (X axis) and treatment (indicated by color). **B: Fraction of pA^+^ reads distribution per read class.** Organized as in panel A, but the Y axis shows the distribution of fraction of pA^+^ reads. **C: Stabilization per read class following RBM7/ZCCHC8 depletion.** Organized as in panel A, but the Y axis shows RBM7 depletion or ZCCHC8 depletion log_2_FC vs Ctrl. To avoid undue influence of outliers, Y axis is capped at log_2_FC=1. **D. Association of genomic/expression features with read class contribution.** Y axis shows selected genomic or expression features. X axis shows Gaussian linear model coefficients relating RNA class proportion (centered log ratio) to each feature separately. Negative values mean a negative correlation between the feature and high contribution of a given read class, and positive values indicate positive correlations (Methods). Bar plots are colored by read class. **E: Stabilization of pA^-^ intermediate and First-intronic class reads following hMTR4/ZFC3H1-depletion.** Organized as in panel C, but the Y axis shows the hMTR4 depletion or ZFC3H1 depletion log_2_FC vs. Ctrl. based on TLDR-seq data. **F: Heat map visualization of nuclear-cytoplasmic ranks of genic reads.** Columns correspond to genes having reads in all classes (see Fig. S2G for all genes). Rows correspond to selected read classes and pA status. Cell color corresponds to nuclear-cytosolic ratio as in panel A. The data shows a split of genes according to whether their First-exonic reads were nuclear or cytosolic enriched (indicated by dotted lines and analyzed further in panel G). **G: Stabilization of First-exonic reads following RBM7/ZCCHC8 depletion, separated by nuclear enrichment.** Organized as in panel C, but only plots First-exonic class reads, which are separated between nuclear and cytosolic-enriched cases. **H: Linear model of RBM7-depletion sensitivity for First-exonic, First-intronic and PROMPT reads.** The X and Y axes are organized as in Fig. 2D, but the linear model was expanded to also include First-exonic (only the nuclear subset) and First-intronic read classes. Each dot is a region, colored by type. Dotted line indicates the fitted trend line, where grey shows the 95% confidence interval of the fit. Density plots on top and to the right show the log_2_FC vs. Ctrl (right) and A/T content distributions for each region type (top). **I: Proposed model for biogenesis and fate of read class transcripts.** See main text for details.

As expected, Full-length reads were predominantly cytosolic, pA^+^ and insensitive to NEXT depletions (Fig.4A-C), consistent with mature mRNA transcript characteristics. Genes enriched for Full-length reads were on average highly expressed and characterized by compact architectures, including genes and transcripts with fewer and shorter exons and introns, and associated with GC-rich isochores, long RNA half-lives and high codon adaptation index values (Fig.4D).

In contrast, Intermediate reads were predominantly pA^-^, cytosol-enriched (Fig.4A-B), and overall, not affected by NEXT and PAXT depletions (Fig.4C,E). Intermediate reads were almost invariably spliced (98.0% of reads) and strongly enriched for 3′ ends within annotated exons (89.2% of reads) where the large majority of 3’ ends (77%) were located within exon bodies and did not overlap splice sites. Genes enriched for Intermediate reads exhibited largely opposite properties to Full-length-enriched genes, including longer genes and transcripts, more and longer exons, less optimal codon usage, and no substantial enrichment of prolonged cytosolic half-lives (Fig. 4D). The increased fraction of Intermediate reads in longer RNAs is unlikely to reflect incomplete second-strand synthesis, as we required detection of both 5’ and 3’ TLDR-seq adapters for read inclusion; importantly, the 3′ adapter is ligated directly to the RNA. We tested whether fragmentation products with phosphorylated 3′ ends contributed to Intermediate-class read fractions by omitting the 3′-end dephosphorylation step in TLDR-seq. This did not decrease the mean per-gene fraction of intermediate reads, arguing against a substantial contribution from dephosphorylation-dependent capture of these products (Fig. S2D). In addition, analysis of previously published QuantSeq 3’ end sequencing data from the same cells, generated with and without poly(A) polymerase (PAP) treatment to polyadenylate and thereby be able to capture polyA^-^ RNAs^24^, further revealed higher fractions of pA^−^ 3′ ends within exon bodies in genes with higher TLDR-seq Intermediate-read fractions, with these 3’ ends distributed broadly across exon bodies, independently supporting the TLDR-seq observations (Fig. S2E). Together, these observations support a model in which Intermediate reads represent capped products of progressive cytosolic 3′→5′ degradation of mature mRNAs. Cytosolic endonucleolytic cleavage events could contribute but is less likely since 3’ ends were typically evenly spread across exons. Consistent with distinct cytosolic fates, Full-length- and Intermediate-enriched genes also had distinct 3′-UTR RNA binding motif profiles (Fig. S2F).

In contrast to these predominantly cytosolic classes, First-intronic reads were highly nuclear-enriched, largely pA^-^ and sensitive to RBM7- and ZCCHC8 depletions (Fig.4A-C). Their abundance was associated with long first introns, short first exons and increased gene length, features consistent with previously described PTT-derived RNAs which are nuclear exosome substrates^11,25–27^. As with PROMPTs, stabilization frequently coincided with 3’end polyadenylation (Fig.S2G).

Genes enriched for First-exonic reads were characterized by long first exons, G/C-rich promoter-proximal regions and elevated evolutionary conservation of promoters and early gene regions (Fig.4D). Although the class showed overall limited sensitivity to NEXT and PAXT perturbations (Fig.4C,E), compartment enrichment analysis revealed nuclear and cytosolic subclasses (Fig.4F, Fig. S2H). We found that there was a higher correlation between the first and remaining exons in terms of reads per nucleotide in the cytosolic subclass than in the nuclear subclass (Fig.S4I). Therefore, cytosolic First-exonic reads most likely represent an extension of the degradation pathway inferred for Intermediate reads, whereas nuclear First-exonic reads are more consistent with PTT events. Given this, we were surprised that nuclear-enriched First-exonic reads were not NEXT sensitive (Fig.4G). We hypothesized that low A/T content in first exons may lead to lower NEXT sensitivity and therefore extended the regression model used to relate PROMPT A/T content and transcript length to NEXT sensitivity (Fig.2D) to include PROMPTs, nuclear First-exonic and (all) First-intronic reads. This showed that once length effects were regressed out, reduced NEXT sensitivity of First-exonic transcripts could largely be explained by their lower A/T content (Fig.4H, S2J).

Together, these analyses identify two major RNA-fate pathways underlying read-class composition at mRNA loci (Fig.4I). In this model, full-length and Intermediate reads predominantly derive from mature, pA^+^ mRNAs that are exported to the cytosol, with Intermediate reads representing progressive 3′→5′ cytosolic degradation products. The cytosolic subset of First-exonic reads likely follows the same pathway. In contrast, First-intronic reads and a nuclear subset of First-exonic reads arise from PTT and subsequent nuclear surveillance. Notably, the different NEXT sensitivities of these prematurely terminated transcript classes can largely be explained by the same transcript-length and sequence-composition features that govern PROMPT degradation.

### Convergent enhancer transcription marks a distinct poised enhancer architecture

Finally, we investigated how eRNA architecture relates to enhancer activity. To avoid defining enhancers based on eRNA TSS configurations, we identified non-exonic DHSs and classified a subset of them as enhancers using orthogonal MGI annotation^29^ based on mESC chromatin and functional data. Restricting the analysis to enhancer regions overlapping >= 10 TLDR-seq 5′ ends yielded 7,704 transcribed enhancers using RBM7-depleted libraries. eRNAs were, like PROMPTs, generally short, unspliced and stabilized upon NEXT depletion(Fig.S3A-C). Stabilized eRNAs were frequently polyadenylated and shorter eRNAs were less strongly stabilized by NEXT depletion than longer eRNAs(Fig.S3D-F).

We first examined whether eRNA length was associated with enhancer activity. Using the 75th percentile of eRNA lengths as a representative length measure, we found that eRNA lengths on opposite strands were not correlated in most cases (Fig. 5A). mESC STARR-seq activity from ref.^30^ increased with representative eRNA lengths (across both strands) up to∼620 nt but declined for longer transcripts (Fig.5B), suggesting that enhancer activity is maximal at intermediate eRNA lengths.

**Figure 5:**
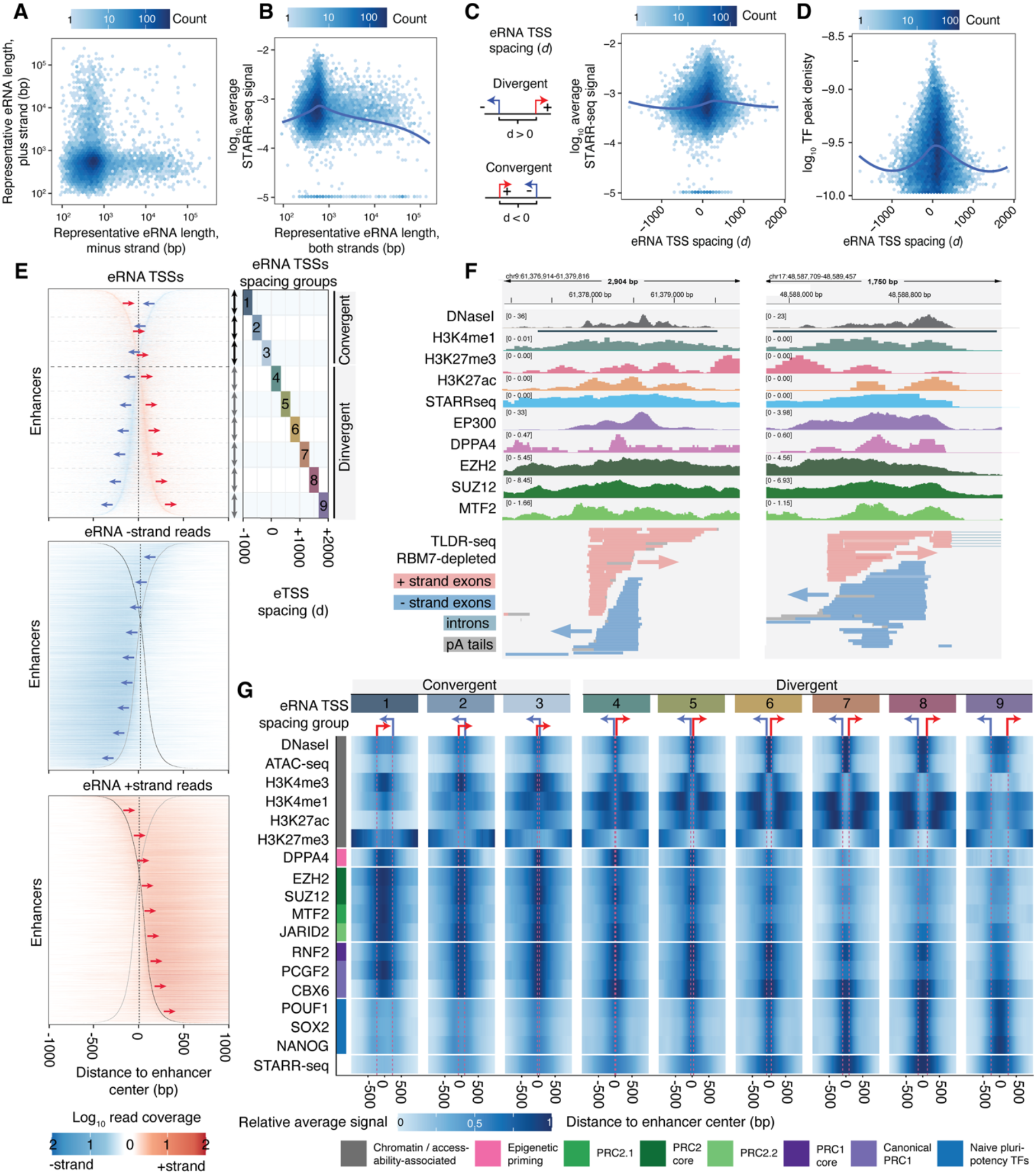
Convergent enhancer transcription is associated with a poised-like chromatin state. **A: Scatter plot of representative eRNA lengths on opposite strands of each enhancer.** Axes show representative eRNA lengths per enhancer strand (75th percentile of TLDR-seq read lengths), log_10_-scaled. Hexagon color represents the number of observations. **B: Relation between enhancer potential and eRNA length.** The plot is organized as in panel A but the X axis shows representative eRNA length on each strand and the Y axis shows the average STARR-seq signal for the enhancer region in log_10_ scale. Trend line in deep blue. **C: Relation between eRNA TSS spacing and STARR-seq signal.** The plot is organized as in panel B but **t**he X-axis shows eRNA TSS spacing (*d*), defined as the plus strand TSS coordinate subtracted by the minus strand TSS coordinate (also see schematic to the left). **D: Relation between eRNA TSS spacing and TF ChIP-seq peak density.** The plot is organized as in panel B but Y axis shows log_10_-scaled ChIP seq binding peaks per enhancer nucleotide based on ReMap data. **E: eRNA read coverage of convergent and divergent bidirectionally transcribed enhancers.** In each heat map, each row is an enhancer region, aligned by the midpoint between dominant eRNA TLDR-seq TSSs (dotted line), and ordered by the eRNA TSS spacing (*d*) as defined in panel C, where convergently transcribed enhancers have negative *d* values. All subplots use the same row order. The X axis shows distance in bp from the enhancer midpoint. Color and arrows indicate TLDR-seq strand, color intensity indicate log_10_ TLDR-seq read coverage. Top heat map shows coverage of 5’ ends of TLDR-seq reads (first 20 nt of reads are plotted). Dotted horizontal lines, and the right side of the plot shows classification of enhancer regions into 9 equally large groups based on *d* (X-axis). The middle and bottom heat map show full, strand-TLDR-seq read coverage on minus or plus strand, respectively. Positions of the most prominent TSSs on both strands are shown as grey lines in both plots. **F: Representative examples of two convergently transcribed enhancer regions**. Plots are organized as in Fig.1A (lower panel) but show additional ChIP-seq tracks. **G: Chromatin and DNA binding protein enrichments around enhancer midpoints as a function of eRNA TSS configuration.** Heat map rows correspond to DNase I-seq, ATAC-seq, ChIP-seq and STARR-seq assays as indicated on the left: see color legend for larger groupings. X axis shows location relative to enhancer midpoint in bp. Each subplot corresponds to an enhancer group as defined in panel E based on eRNA TSS configuration and spacing: dotted lines show average TSS locations within each group, schematic on top shows TSS configuration and strand. Heat map cell color corresponds to the average genomic assay read depth across all enhancers in the group and then normalized so that min=0, max =1 across the row.

We next analyzed the relationship between enhancer activity and eRNA TSS spacing, defining *d* as the signed genomic distance between the dominant plus and minus-strand eRNA TSSs. Positive *d* values correspond to divergent TSS pairs, whereas negative values correspond to convergent pairs (Fig.5C, left). STARR-seq signal increased with *d* at small positive *d* values, plateaued when *d* was between ∼+250 to +500 bp and declined at larger separations (Fig. 5C, right). Because transcription factor (TF) occupancy is a major determinant of enhancer activity, we quantified mESC TF ChIP-seq peak density within each enhancer using ReMap^31^ data. Although STARR-seq signal increased with TF occupancy (Fig.S3G), TF density was highest at the smallest TSS positive *d* values and declined with increasing *d* (Fig.5D), indicating that the relationship between TSS spacing and enhancer activity cannot be explained by TF density alone.

Unexpectedly, ∼33% of transcribed enhancers exhibited negative *d* values, corresponding to convergent eRNA transcription (Fig.5E, left, examples in Fig.5F). Importantly, STARR-seq activity was maximal between the convergent eRNA TSSs (Fig.S3H). To determine whether convergent enhancers represent a distinct regulatory state, we stratified enhancers into nine equally sized groups by eRNA TSS spacing (based on *d* values: Fig.5E, right) and examined TF and chromatin-factor occupancy using mESC ChIP-seq data from ReMap^31^. Visualization of average ChIP-seq signal centered on enhancer midpoints (Fig.5G) showed that divergently transcribed enhancers displayed the expected active enhancer architecture, with a central nucleosome-depleted region (NDR), flanking enhancer-associated histone modifications (H3K4me1 and H3K27ac), and strong TF occupancy within the NDR. Convergent enhancers showed a markedly different organization. Their central regions were less accessible and enriched for PRC2 components, including EZH2, SUZ12, MTF2 and JARID2, together with histone modifications, most prominently H3K4me3 and H3K4me1, but also more weakly H3K27ac and H3K27me3. H3K27me3 enrichment was stronger further outside the enhancer center, a pattern more pronounced at convergently vs. divergently transcribed enhancers. Together, the coexistence of enhancer-activity-associated histone modifications with PRC2 occupancy and H3K27me3 suggested a poised or bivalent-like chromatin configuration, consistent with previous descriptions of Polycomb-associated poised enhancers in mESCs^32,33^.

Most TF occupancy closely followed accessibility and was reduced at convergent enhancers, including pluripotency factors (e.g. POU5F1, SOX2, NANOG) that were strongly enriched within the NDR of divergent enhancers. A striking exception was DPPA4, which was selectively enriched in the convergent classes and closely followed the PRC2 occupancy pattern. This is notable given the established role of DPPA2/4 as epigenetic priming factors at PRC-associated, H3K4me3/H3K27me3-bivalent promoters in mESCs^34^, supporting an association between convergent enhancer architecture and a poised chromatin state. PRC1 protein occupancy was comparatively invariant across enhancer architectures. STARR-seq signal was lowest at convergent enhancers, but as also noted above (Fig. S3H), a localized activity maximum was retained at their centers.

We next asked how transcription initiation is organized within this distinct chromatin architecture. Because enhancer-centered averaging blurs features associated with individual TSSs when TSS spacing varies between loci, we performed an eRNA-TSS-centered analysis by anchoring the profiles on the plus-strand eRNA TSS of each enhancer, focusing on convergent enhancers with the largest eRNA TSS spacing (group 1), using a divergent enhancer class for comparison (group 7; Fig.6A).

**Figure 6:**
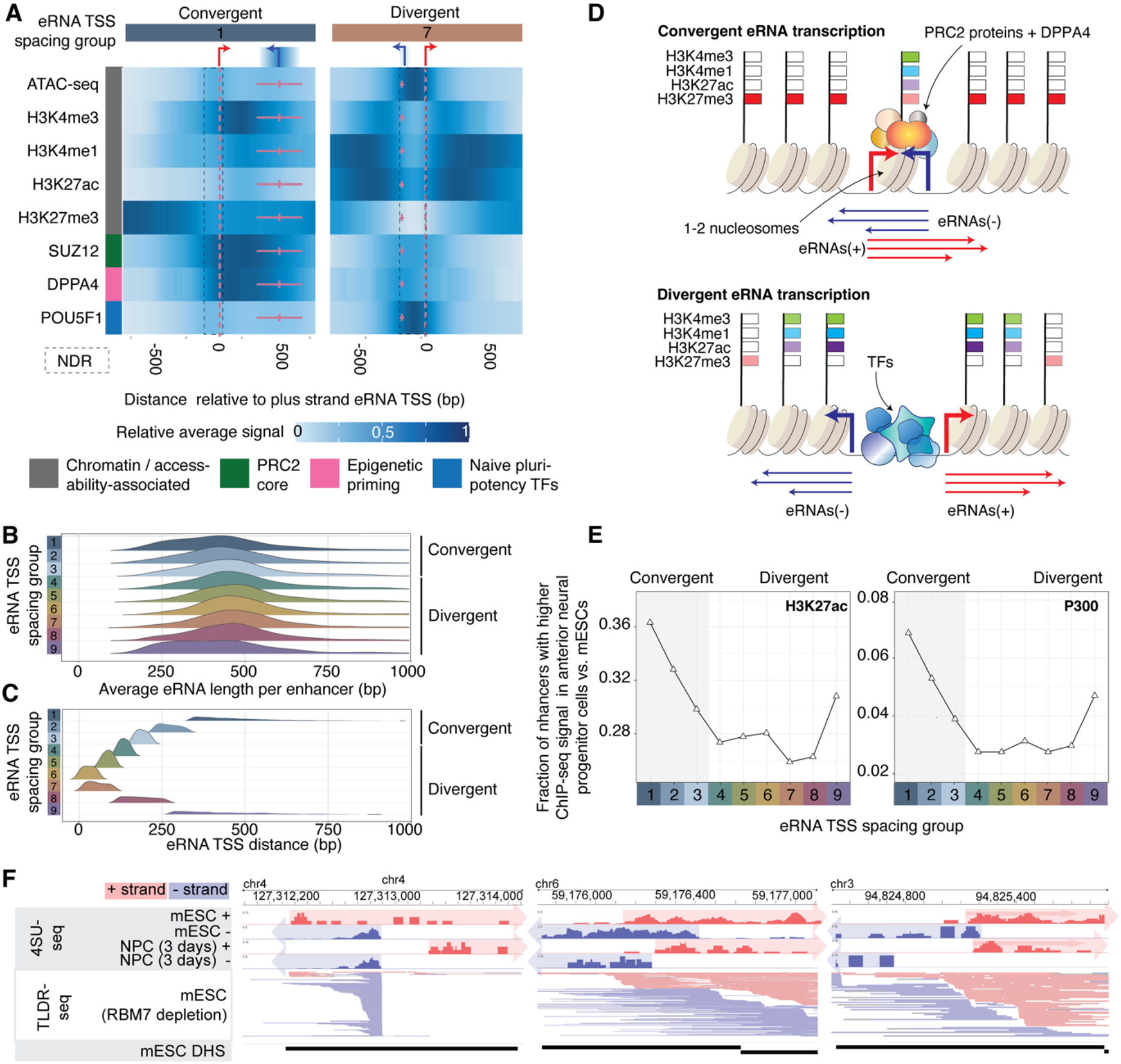
TSS positioning and activation potential of convergently transcribed enhancers. **A: Chromatin and DNA binding protein enrichments around enhancer midpoints as a function of eRNA TSS configuration.** Heat map is organized as in Fig. 5G, but aligned at dominant eRNA plus strand TSSs of each enhancer region (x=0). Only enhancer groups 1 (highly convergent eRNA TSSs) and 7 (typical divergent eRNA TSSs) and selected data rows are shown. Average locations of the dominant minus strand eRNA TSS are shown as a horizontal pink line, while its corresponding standard deviation is shown as a horizontal line. Black dotted lines indicate the inferred boundary of the nucleosome-depleted region discussed in the main text, based on average ATAC-seq and histone modification ChIp-seq signals. **B: Distribution of eRNAs lengths per enhancer group.** The ridge density plots show the distribution of average eRNA lengths (X axis) per eRNA TSS spacing group as defined in Fig. 5E (Y axis). Fill colors correspond to eRNA TSS spacing group. **C: Distribution of eRNA TSS distance per enhancer group.** Organized as panel B but X axis shows absolute distance in bp between eRNA TSSs (same scale as in B). Fill colors as in B **D: Schematic model showing overall properties of convergently vs. divergently transcribed enhancers.** See Main text for discussion. **E: Propensity for gaining active enhancer features after differentiation to anterior neural progenitor cells as a function of eRNA TSS configuration.** The Y-axis shows the fraction of enhancers that have higher ChIP-seq signal in anterior neural progenitor cells vs. mES cells (log_2_FC>0 in the +-250bp region around each enhancer midpoint). X axis shows eRNA spacing group as defined Fig 5E. Each panel shows one ChIP-seq analysis; P300 and H3K27ac ChIP-seq data from ref^33^. **F: Examples of convergently transcribed eRNA configurations shifting to divergent configurations as a function of differentiation.** Genome-browser examples show 4sU-seq read density at three enhancer locations at naive mESC and after 3 days of differentiation towards neural progenitor cells (NPCs), colored by read strand. Arrows behind the data show predicted TSS and eRNA body (see Methods). TLDR-seq data and DHS locations from naive mESCs are shown below.

As expected, eRNA TSSs at divergently transcribed enhancers flanked the central NDR and initiated transcription outwards. However, at convergently transcribed enhancers, the eRNA TSSs were positioned at the boundary of the less accessible central region, close to the inner edge of a short local NDRs flanking it (also visible in 5G), and initiated transcription inwards towards the enhancer center. Notably, the central, less accessible central region downstream of the eRNA TSSs had increased H3K4me1, H3K4me3, H3K27ac, H3K4me3, PRC2-component and DPPA4 signal, whereas other TFs, exemplified by POU5F1, showing only modest enrichment and preferentially occupied the local NDR upstream of the eRNA TSS. Because eRNA length distributions were largely invariant across TSS-spacing groups, with median lengths of approximately 375 nt and only modest shortening at the largest TSS separations (Fig.6B), eRNA transcription length did not appear to be strongly constrained by either the downstream nucleosome boundary or the opposing TSS (Fig.6B,C). Despite these striking differences in TSS positioning between enhancer architectures, we found relatively little sequence information that could explain eRNA TSS positioning (Fig.S4A), suggesting that local chromatin architecture may play a greater role in this process. Thus, our data support a model in which transcription at convergently transcribed enhancers initiates from the edges of a central PRC2-associated nucleosome with a poised or bivalent-like chromatin configuration (Fig.6D).

We reasoned that if convergent enhancer architecture marks poised enhancers, some of these loci may become activated during differentiation from mESCs to other cell states. Importantly, we would only expect a subset of convergently transcribed enhancers in mESCs to become active in a given differentiation state: some poised enhancers may remain poised to be activated in other states, or never become active. We therefore examined changes in chromatin and DNA-binding protein occupancy across a diverse collection of mESC differentiation datasets from ReMap^31^. The clearest relationship was observed during differentiation towards anterior neural progenitor cells (AntNPCs)^33^, where increasingly convergent enhancer architectures in mESCs were associated with increased p300 binding and H3K27ac ChIp seq in AntNPCs vs. mESCs, indicative of enhancer activation in AntNPCs (Fig.6E). Concordantly, using nascent RNA data (4sU-seq and GRO-seq) from two separate naive mESC to neural progenitor cell (NPCs) differentiation experiments^35,36^, we found that 12% (4sU-seq, 58/464) and 11% (GRO-seq, 89/781) of convergently transcribed enhancers in mESCs, as defined by respective data set, became divergently transcribed in NPCs (see examples in Fig.6F). These observations are consistent with convergently transcribed enhancers representing a poised enhancer state with the potential for activation, and divergent eRNA transcription, during neural differentiation or other developmental transitions.

## Discussion

Here, using ultra-deep sequencing of capped, full-length pA^+/−^ RNAs, we linked transcript architecture to RNA fate at promoters, genes and enhancers. This revealed length- and sequence-associated PROMPT sensitivity, abundant capped cytosolic mRNA degradation intermediates and a Polycomb-associated poised enhancer architecture.

PROMPT polyadenylation scaled with transcript stabilization, was enriched among longer transcripts and lacked canonical polyadenylation-site features, supporting a post-stabilization rather than cleavage-coupled polyadenylation^19^. Independently, transcript length and local A/T content predicted NEXT sensitivity, implicating both architecture and sequence context in nuclear exosome targeting.

Surprisingly, intact mRNAs constituted a minority of the capped protein-coding RNA molecules detected by TLDR-seq. Absolute read-class proportions should, however, be interpreted cautiously, as they may be influenced by technical biases, including preferential sequencing of shorter molecules by nanopore sequencing. The key observation is therefore not the precise fraction of reads or genes assigned to each class, but the pronounced differences in read-class composition between genes, which in turn were strongly associated with gene architecture and function: Full-length transcripts predominated among highly expressed, compact genes, whereas Intermediate transcripts were enriched among longer, exon-rich genes. Nuclear First-exonic and First-intronic classes were consistent with promoter-proximal termination and exosome surveillance, governed by the same length- and sequence-related NEXT sensitivities as PROMPTs. The strong coupling between read-class composition, gene architecture and gene function raises the possibility that different transcript architectures reflect distinct expression strategies. Although gene architecture, expression level and sequence composition were strongly associated with read-class composition, they do not fully explain why particular genes are enriched for specific read classes, especially the widespread Intermediate class. It also remains unclear whether read-class composition is stable across cell types and states. In addition, because TLDR-seq captures capped RNAs only, the relative contributions of capped versus uncapped intermediates generated by 5′→3′ decay remain incompletely resolved.

Our enhancer analyses suggest that eRNA TSS positioning captures information about enhancer state: ∼33% of transcribed enhancers had convergent TSSs, PRC2 occupancy and poised chromatin features. Consistent with this, a subset of convergently transcribed enhancers exhibited evidence of activation during differentiation towards neural progenitor cells, as evidenced by ChIP and nascent eRNA data. Thus, transcriptionally active enhancers can coexist with Polycomb-associated poised chromatin, while their transcriptional architecture conveys regulatory information beyond total eRNA output. An important question is whether convergent eRNA transcription contributes directly to enhancer state or primarily reflects the underlying chromatin architecture. Although our observations favor the latter interpretation, previous studies have reported functional roles for enhancer transcription and eRNAs, including links to Polycomb regulation, as reviewed by^37–40^.

Together, our analyses show that full-length transcript architecture links RNA fate to genome regulation across promoters, genes and enhancers.

## Methods

### Cells and exosome factor depletions

All mESC lines used or generated in this study were descendants of wildtype ES-E14TG2a cells (male genotype, XY). mESCs were cultured on 0.2% gelatin coated plates in 2i/LIF containing media (1:1 mix of DMEM/F12 (Thermo Fisher, #31331-028) and Neurobasal (Thermo Fisher, #12348-017) supplemented with 1% (v/v) penicillin-streptomycin (Sigma, #P4333), 2 mM Glutamax (Thermo Fisher, #35050-038), 0.1 mM Non-Essential Amino Acids (Thermo Fisher, #11140-035), 1 mM Sodium Pyruvate (Thermo Fisher, #11360-039), 0.5x N-2 Supplement (Thermo Fisher, #17502-048), 0.5x B-27 Supplement (Thermo Fisher, #17504-044), 50 µM 2-Mercaptoethanol (Thermo Fisher, #31350-038), 3 µM GSK3-inhibitor (CHIR99021, Sigma, #SML1046), 1 µM MEK-inhibitor (PD0325901, Sigma, #PZ0162) and Leukemia Inhibitory Factor (LIF, produced in house in the THJ group)) at 37°C, 5% CO₂. Cells were passaged every 48-72h by dissociating cells with 0.05% Trypsin (Sigma, #T3924) before neutralizing with an equal volume of 1x Trypsin Inhibitor (Thermo Fisher, #17075-029). Cells were pelleted by centrifugation to remove Trypsin before resuspending in 2i/LIF media and plating at ∼ 8×10^4^ cells/ml.

The generation of CRISPR/Cas9-mediated genomic knock-ins of N-terminal 2xHA-FKBP-V(HA-dTAG)-RBM7 mESCs was described in^41^ and HA-dTAG-ZCCHC8, hMTR4-HA-dTAG were generated in a similar way. The generation of OsTIR1-HA Zfc3h1-3F-mAID mESCs (referred to in text as ZFC3H1-3F-mAID) was described in^24^. Depletion of HA-dTAG tagged proteins was performed by the addition of dTAG^V^-1 (Tocris, #6914) to the cell culture medium for 4 h at a concentration of 100 nM, and wild type (WT) cells with the same treatment were used as controls. Depletion of ZFC3H1-3F-mAID was performed by the addition of indole-3-acetic acid (Sigma, #I5148) for 4 hr at a concentration of 750 µM. ZFC3H1 rapid depletion was complemented by expression of mmZfc3h1 cDNA or an empty vector, and the cell line expressing mmZfc3h1 cDNA was used as the control condition.

Mouse Zfc3h1 cDNA was amplified from a full-length mouse pCR-XL-TOPO[mZFC3H1] cDNA plasmid (Transomic Technologies) using Phusion DNA polymerase (ThermoFisher, #F534). The fragment was cloned using NEBuilder HiFi DNA assembly (NEB, #E2621) into a piggyBAC (’pB’) vector harbouring a C-terminal triple FLAG tag and the BSD gene encoding blasticidin resistance. The construct was fully sequenced. ZFC3H1-3F-mAID mESCs were transfected with the pB[ZFC3H1-3F]BSD plasmid or empty vector control (pB[-3F]BSD) along with a pB transposase expressing vector (pBASE) using Lipofectamine 3000 (Invitrogen, #L3000001). Cell pools were selected using 20 µg/mL blasticidin (Gibco, #R21001) for 7 days. The expression of ZFC3H1-3F was validated by Western blotting using anti-ZFC3H1 antibody following depletion of endogenously tagged ZFC3H1-3F-mAID.

### Nuclear-cytosolic fraction preparation

For nuclear-cytosolic preparations, cells were detached from plates using 0.05 % Trypsin-EDTA (Sigma, #T3924), which was neutralised by addition of 1x Trypsin Inhibitor (Thermo Fisher, #17075-029). Cells were pelleted by centrifugation at 200 x g. All centrifugation steps were performed at 7°C to avoid inadvertent freezing. Cells were washed twice with ice-cold PBS then resuspended in hypotonic buffer (10 mM HEPES, 50 mM KCl, 0.1mM EDTA, 0.1 mM EGTA, 1 mM DTT and 1X protease inhibitor (Roche, #04693159001)) and incubated on ice for 15 minutes. Cells were lysed by addition of 10 % NP-40 to a final concentration of 0.6 % and incubated for a further 5 minutes on ice. Nuclei were pelleted by centrifugation at 1000 x g for 5 minutes. 60 % of the supernatant was removed and underwent additional centrifugation at 16,000 x g, with the resulting supernatant collected as the cytoplasmic fraction and divided for RNA isolation or western blot analysis. Nuclei were resuspended in hypotonic buffer and separated for RNA isolation and western blot analysis. Nuclei were then washed a further two times in hypotonic buffer supplemented with 0.1% NP-40. For western blot analysis, nuclei were lysed in low-SDS TOPEX buffer (300 mM NaCl, 50 mM Tris HCl, 1 mM MgCl2, 1 mM DTT, 0.5 % Triton X-100, 0.1 % SDS, protease inhibitor (Roche, #04693159001) and 250 U/mL benzonase (Sigma, #E1014)) and incubated for 10 minutes at room temperature. The resulting soluble fraction was collected. Western blot lysates were prepared by addition of LDS sample buffer (Invitrogen, #NP0007) and reducing agent (Invitrogen, #NP009) to the cytoplasmic or soluble nuclear fractions. 2 µg protein was loaded for each sample. RNA was isolated by adding TRIzol (Invitrogen, #15596018) directly to pelleted nuclei or in a 4:1 ratio to the cytoplasmic fraction. Nuclear fractions were mechanically homogenized using a 21G needle to break up chromatin. Isolated RNA was DNase treated for 30 minutes using the Turbo DNA-free kit (Invitrogen, #AM1907) before a final clean-up by ethanol precipitation.

### TLDR-seq library preparations and read processing

TLDR-seq libraries were prepared as in^16^. Initial TLDR-seq libraries (6 Ctrl, 3 RBM7- and 3 ZCCHC8-depletions) were prepared using the SQK-LSK110 kit (Oxford Nanopore Technologies) and sequenced for 72 h in R9.4.1 PromethION flow cells (FLO-PRO002), using the software MinKNOW. Nuclear and cytosolic RNA fractions of wild type samples were prepared using the SQK-LSK114 kit (Oxford Nanopore Technologies) and sequenced in an additional PromethION R10.4.1 flow cell (FLO-PRO114) using the same computational pipeline. ZFC3H1, hMTR4 and Ctrl complementary TLDR-seq libraries were prepared using the SQK-LSK110 kit (Oxford Nanopore Technologies) and sequenced in a R9.4.1 MinION flow cell (FLO-MIN106D). Reads were basecalled using Dorado 0.7.1 with “--no-trim”, “--estimate-poly-a”, and “sup” options. Poly(A) tail lengths were estimated directly from the raw Nanopore squiggle files using a custom C++ algorithm integrated into the dorado basecalling pipeline, adapted from the tailfindR^42^ method. For each transcript, raw squiggle arrays were Z-normalized, rectified to absolute values, and truncated to handle signal spikes. To isolate the characteristic low-variance electrical signature of poly(A) tails, the normalized signal was smoothed using a sliding-window moving average approach. Poly(A) boundaries were defined by scanning for continuous regions with near-zero local slopes and low electrical variance relative to the adjacent transcript body. Final poly(A) tail lengths in nucleotides were deduced by dividing the total raw signal duration of this low-variance region by the dynamic estimate of signal samples per nucleotide. Basecalled sequencing reads were processed using cutadapt^43^ to remove adapter sequences and sample-specific barcodes. Stringent two-sided trimming was enforced, requiring the explicit detection of both 5’ and 3’ adapter structures, and untrimmed reads were systematically discarded. The resulting curated reads were aligned against the mouse reference genome (GRCm39/mm39^44^) using minimap2^45^ with splicing-aware parameters (-ax splice). To resolve multi-mapping ambiguities and retain high-confidence genomic loci, secondary and unmapped alignments were systematically excluded using samtools. For sequencing reads exhibiting multiple primary alignments, a custom parsing algorithm was applied to isolate and retain strictly the single alignment harboring the maximum alignment score. To account for the directionality of the library preparation chemistry, alignments were properly restranded to reflect the true transcriptomic strand of origin, using the adapters sequence strand information. Finally, the strand-corrected, uniquely assigned alignments were demultiplexed according to their sample barcodes.

### Definition of DHS-anchored TLDR-seq clusters and initial analyses

Publicly available DNase-seq data were downloaded from the ENCODE portal^17^ (accession ENCSR000CMW). Fastq files of technical replicates were concatenated into two replicate pools and aligned against the mouse reference genome (GRCm39/mm39) using bwa mem^46^. Initial DNase I peaks were called using MACS3^47^ (callpeak) with a strict false discovery rate (q-value) threshold of 0.05 to generate a preprocessed set of hypersensitive regions and corresponding genome-wide coverage tracks. To refine DHS boundaries and resolve multimodal hypersensitive regions, we decomposed the peaks using a custom signal algorithm. First, the DNase I continuous coverage was tiled into 50-bp genomic bins and smoothed using a running mean across a 100-bin window. Then, the initial MACS3 peaks were extended by 600 bp in both directions, and overlapping peaks were merged to capture broad regulatory zones. DNase I sequencing depth was then calculated across these extended peaks to identify local signal maxima (summits) and their flanking local minima. The summits were filtered according to 3 criteria: a valid summit had to 1) reach at least 30% of the extended peak global maximum signal, 2) be at least 1.6 times higher than the adjacent minima, and 3) be separated from adjacent summits by a minimum distance of 250 bp. The final boundaries for each decomposed DHS were determined by the genomic coordinates of the flanking local minima.

Genomic TLDR-seq alignments were assigned to the above-defined DHSs using bedtools^48^, requiring the 5’ start coordinate of each read to fall within the boundaries of a given DHS. TLDR-seq reads on each strand then formed DHS-anchored, strand-specific TLDR-seq clusters. We normalized the DHS read counts using scaling factors calculated from the mitochondrial read coverage of each respective library. Principal component analysis on log-transformed normalized count matrices was used to assess global sample reproducibility. Cluster-assigned reads were annotated with relevant mapping features, including genomic coordinates, exon counts, and strand orientation. Transcripts were additionally stratified by their polyadenylation status (classifying reads with poly(A) tails strictly greater than 15 nucleotides as polyadenylated) and partitioned into discrete length bins (ranging from 0-100 bp to >1500 bp) based on their total genomic span to evaluate length-dependent dynamics. To identify TLDR-seq clusters with altered read abundance upon exosome-related factor depletion, differential expression testing was performed using DESeq2^49^, comparing Ctrl samples against each targeted depletion condition (RBM7- and ZCCHC8 depletions). Significantly upregulated clusters were defined using a log_2_FC > 1 and *FDR* < 0.05 threshold. When relevant, we remade the same analysis but stratified reads from each cluster by length (e.g. Fig. 2B).

To assess the structural signatures of transcripts accumulating upon exosome-factor depletion, targeted structural comparisons were restricted strictly to transcripts originating from DHS regions identified as significantly upregulated in the targeted knockdown conditions relative to Ctrl. For these specific subsets, mean and median values for exonic length, exon count, genomic span, and poly(A) tail length were computed. Splicing complexity was evaluated by categorizing these transcripts by exon count and comparing their fractional distributions across genotypes.

### Definition of PROMPT TLDR-seq clusters

To identify PROMPT TLDR-seq clusters, we identified first transcribed DHSs that overlapped RefSeq-annotated mRNA TSSs but not annotated transcription termination sites or annotated lncRNAs and then extracted the TLDR-seq cluster from these DHS that was on the opposite strand of the mRNA.

### PROMPT polyadenylation analysis

PROMPT reads were anchored to their 3’ ends, excluding any reads overlapping downstream protein-coding genes. Local termination maxima were identified using Kernel Density Estimation to call cleavage “peaks” independently for pA^+^ and pA^-^ populations. Shifts between pA^+^ and pA^-^ termination coordinates within the same expressed PROMPT were statistically evaluated using Wilcoxon Rank Sum tests. Genomic sequences flanking the curated 3’ peaks were extracted. *De novo* motif discovery was performed utilizing XSTREME^50^, contrasting regions targeted by specific knockdown conditions against non-targeted or Ctrl backgrounds. Discovered position-specific weight matrices of putative RNA-binding factors were mapped back to the extracted sequences to evaluate their positional frequency and spatial distribution relative to the mapped cleavage sites. To correlate degradation sensitivity with polyadenylation status, PROMPT TLDR-Seq clusters were stratified by their differential expression levels using k-means clustering and variations in the polyadenylated read fraction across intervals were statistically tested by Analysis of Variance (ANOVA).

### Modeling of PROMPT stabilization following exosome depletions as a function of length and sequence composition

To investigate the influence of sequence composition on transcript sensitivity to the nuclear exosome, for each PROMPT, we identified its main TSSs and defined consecutive 20 bp windows starting from this TSS and continuing downstream. In each such window we calculated fraction of A/T nucleotides and the depletion vs Ctrl log_2_FC using TLDR-seq reads whose 3’ ends were located within the window. To decouple the influence of transcript length from the A/T nucleotides we modeled the log_2_FC as a combination of the two effects, using the linear model

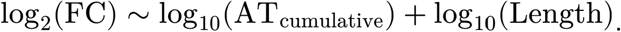

The isolated effect of transcript length (derived from the model’s length coefficient) was regressed out from the raw differential expression values. This yielded a length-corrected log_2_FC metric, allowing for the direct correlation of exosome sensitivity with cumulative AT counts.

### Gene-centric analysis and classification of TLDR-seq reads

To identify genic TLDR-seq clusters, we identified transcribed DHSs that overlapped with RefSeq-annotated mRNA TSSs. Only annotated multi-exonic RefSeq mRNAs were analyzed, and we only analyzed cases where there was a one-to-one match between DHS and annotated TSS per gene (so, genes with two or more highly distinct expressed alternative promoters were not analyzed). For each DHS-gene pair we extracted the TLDR-seq cluster that were on the same strand as the linked gene.

We defined the representative transcription termination site (TTS) of each gene via the peak of the denoised, smoothed poly(A)+ WT 3’-end depth profile using a Gaussian kernel (bandwidth 30 bp). Where the depth profile called a single candidate peak, it was taken as the TTS. Where multiple candidates were called, we selected the representative TTS that met at least two of three criteria (in priority order): closest to RefSeq Select annotated 3’ end, greatest local read support, and most 3’-extended in the direction of transcription. Where all three criteria pointed to different candidates, the read supported candidate was taken. Where no candidate could be called (i.e. genes with low pA+ read counts or flat 3’ depth profile), the RefSeq Select annotated 3’ end was used as TTS.

To classify 3’ end locations, we calculated the distance from the 3’ end of each read to the representative TTS. Transcripts were sorted into one of these distinct structural categories based on the following: Full length: The read 3’ end was within the +-50 bp region around the annotated TTS. First-exonic: The read was unspliced and its 3’ end was located before the boundary of the first annotated exon. First-intronic: The read was unspliced and its 3’ end located within the first annotated intron. Intermediate: 3’ ends were located after the annotated first intron and before the last genomic protein-coding nucleotides (including introns) and not assigned to other classes. We chose this definition to avoid ambiguous cases between classes. Finally, reads were split by polyadenylation status: those with a poly(A) tail longer than 15 nucleotides were assigned pA^+^, otherwise pA^-^.

### QuantSeq analysis of Intermediate reads

We reanalyzed QuantSeq 3′ mRNA-seq FWD + UMI (Lexogen) libraries from wild-type mESCs^24^ (GEO accession GSE178550). Each of three RNA samples was split into an untreated half (noPAP) and a half polyadenylated *in vitro* with *E. coli* poly(A) polymerase (EPAP). ERCC spike-ins (ThermoFisher, ERCC Mix1 #4456740) were added to every library. Cutadapt removed the first 10 nt of each read (6-nt UMI and spacer), trimmed low-quality bases (*q* < 20) and 3′ poly(A) stretches (≥ 8*A*), and discarded reads shorter than 20 nt. Reads were aligned to mm39 with STAR^51^ (uniquely mapped reads only, allowing soft clipping). Reads with identical UMI, and whose 5’ ends mapped to the same location, were collapsed.

The 3′ end of a read without poly(A) was predicted (*m* − 1)*nt* downstream of the read 5′ ends, where *m* is the median fragment length of reads without poly(A), measured in each library at isolated TTS (123 − 125*nt* in noPAP and 135 − 136*nt* in EPAP libraries). For reads containing poly(A), the 3′ end was the last aligned base. For each gene we used the Refseq transcript annotation used in the previous section. 3’ ends in the first exon or in any intron were discarded, as were ends more than 1,050nt past the TTS. The remaining ends were assigned to the region from the start of exon 2 to the CDS end (spliced; 100 bins of equal length), or to the 3′ end of the gene (CDS end to TTS + 1,050 nt). We kept multi-exon genes detected by TLDR-seq whose CDS ended after exon 1 and with ≥ 10 reads in the denominator (n = 3,411). For each noPAP/EPAP pair *i*, the scale factor *s_i_* was the ratio of EPAP to noPAP ERCC counts (0.53, 0.62, and 0.38). Within each genomic bin, non-polyadenylated pA^-^ ends were estimated as *EPAP_i_ - s_i_ · noPAP_i_*. In noPAP, ends up to the CDS end come from polyadenylated RNA or from oligo(dT) priming on genomic A-rich sequence (internal priming). Assuming internal priming scales like the rest of the library between noPAP and EPAP, the subtraction removes it. All EPAP ends at the 3′ end of the gene were considered biological. For each gene *g* and bin *n*, the cumulative share of pA^-^ ends was:

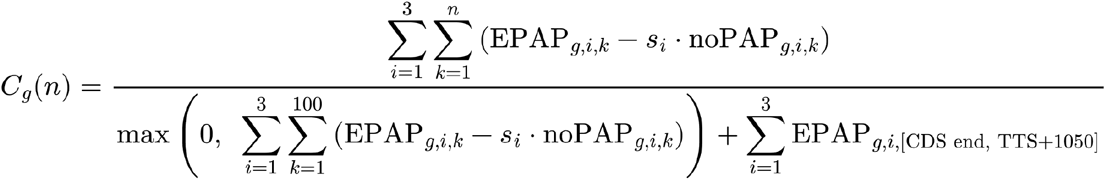

Genes were grouped by their TLDR-seq intermediate pA^-^ fraction, computed over the same read species as here (< 30%, *n* = 991; 30 − 80%, *n* = 1,909; > 80%, *n* = 511). We plotted the mean *C_g_(n)* per group ± s.e.m.

### Nuclear-cytosolic ratio calculation

TLDR-seq reads from cytosolic and nuclear mESC compartments from WT cells (3 replicates each) were processed as previous libraries, but TLDR-seq clusters were normalized using counts per million (CPM) in each library and then averaged across replicates. Next, we calculated a nuclear-to-cytosolic ratio for every TLDR-seq cluster, adding a pseudocount of 1:

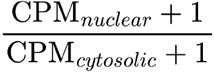

We then used this ratio to sort TLDR-seq clusters by the cytosolic vs. nuclear preference.

### Generalizing the A/T PROMPT stabilization models to all nuclear products

We extended the sliding-window stabilization modeling described for PROMPTs (see section *Modeling of PROMPT stabilization following exosome depletions as a function of length and sequence composition)* to other nuclear species, *i.e.* all first-intron termination events and nuclear first-exon termination products. Consecutive 20 bp windows were generated downstream of dominant TSSs for each class, and any windows physically overlapping annotated splice junctions were masked to avoid sequence biases. To assess whether sequence-dependent sensitivity follows a universal, non-linear trajectory across all distinct read classes (PROMPTs, First-intronic, and nuclear First-exonic), a unified second-degree polynomial regression model was applied to the merged dataset using the formula

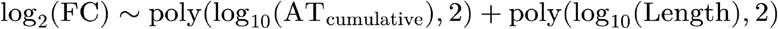

Length-dependent effects were regressed out from the polynomial predictions to isolate the relationship between cumulative AT accumulation and exosome sensitivity (Fig. 4H).

### Feature over/under-representation analysis of read classes

To study genomic and transcriptional features associated with read classes, we conducted Gaussian regression analysis based on Ctrl cell reads, on 4,688 multi-exon genes with total read counts ≥ 20. To optimize interpretability, we chose a single-predictor analysis instead of a multivariate one. Specifically, for each feature and each read class, one single-predictor regression was conducted, associating centered log ratio of read class with the targeted feature:

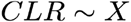

Before regression, feature data was standardized:

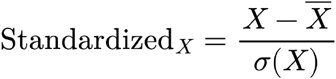

The following candidate features were selected for the association analysis, spanning transcription, splicing, decay, translation, context and evolution: 1) Gene expression: using normalized-count mean log_10_(baseMean + 1) via DESeq2^49^. 2) Annotated gene length: using gene length (bp) based on coordinates of TSS/TTS in RefSeq Select reference database (https://www.ncbi.nlm.nih.gov/refseq/refseq_select/). 3) Annotated RNA length: using concatenated exon length (bp) of the RefSeq Select reference. 4) Nuclear / Cytosolic half-life: using published estimates of RNA half-lives by RNA-flow kinetics in mESC from^52^. 5) Codon Adaptation Index: using calculated values via the CAI function of R package coRdon (v1.31.0) for each full RefSeq coding sequence. Values closer to 1 indicating strong adaptation to preferred codons in highly translated genes. We selected 48 expressed ribosomal genes as reference. 6) Using the isochore definitions established by^53^ we binned the genome into 100kb windows and assigned these windows to their corresponding isochore families based on GC content. Genes were assigned to isochores based on genomic overlap. For genes spanning multiple isochores, assignment was determined by the maximum base-pair overlap. 7) G/C or A/T fraction : using average GC% or AT% values based on mm39 reference genome. 8) Conservation: using evolutionary conservation score phastCons across vertebrate species. Scores over specific regions were calculated with gscores function of R package GenomicScores^54^ on phastCons35way.UCSC.mm39 from the UCSC genome browser^55^.

### Motif enrichment in 3’ UTRs

To calculate the RBP density, RNA binding motifs of Mus Musculus were downloaded from CISBP-RNA^28^ build 2.00. We counted the occurrence RBP binding sites (forward and reverse, separately) of each motif on the 3’ UTR region of each gene through FIMO^56^ version 5.5.8 with a default threshold, *p* < 10^-4^. We composed the query motifs by two sets: its motif in CISBP (denoted as forward), and its reverse complement motif (denoted as reverse). The union of the two sets were counted using –norc setting, so that direction sensitive binding can be distinguished rather than aggregated. To define 3’ UTRs, we used the RefSeq select annotation. Subsequently, densities of predicted binding sites were calculated with occurrence / length of 3’ UTR regions. In total, the density of 904 (457 forward, 447 reverse) motifs was calculated.

To quantify the impact of RNA binding proteins (RBPs) on the proportion of RNA classes, we did a regression analysis. Centered log-ratio (CLR) was calculated for each RNA class in Ctrl as:

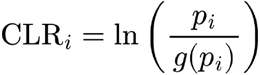

where *p_i_* represents the observed proportion of RNA reads in the total reads for gene *i*, and *g(p)* is the geometric mean of proportions for all possible RNA classes. We regressed the *CLR* of full length RNA (pA^+^) and intermediate (pA^-^) against the density of RBP predicted sites. To control the impact of multicollinearity, we applied a regularization technique, elastic net with a mixing parameter α = 0.5 and λ selected by 10-fold cross-validation using lambda.min. The resulting regression coefficients were extracted for visualization.

### Enhancer curation

To define potential enhancer-DHS, we first overlapped DHS with MGI curated cis-regulatory sites^29^. Enhancers were defined as DHS that fulfilled the following filters: (1) must overlap with an MGI curated enhancer; (2) must not contain RefSeq Select gene TSS, which is defined as a single base coordinate with no flanking window; (3) must be in intronic or intergenic regions. For Figs. 5E-G and 6A-C,D we restricted the analysis to enhancer DHSs exhibiting bidirectional transcription, defined by the presence of both a forward (+) and a reverse (-) strand enhancer TLDR-seq cluster whose 5’ ends overlapped the enhancer DHS. For each DHS, we calculated a DHS midpoint, defined as the exact midpoint between the mean TSS coordinates of the forward and reverse strands. We measured the signed distance *d* between these TSS coordinates (<0 if forward is upstream of reverse, >0 otherwise) and stratified DHSs into 9 quantile groups based on this distance. We then extracted RBM7-depleted library reads whose 5’ ends overlapped each DHSs.

### Processing of ATAC-Seq and histone modification ChIP-seq

We used published ATAC-Seq and histone modification ChIP-seq data generated in mESCs with similar culture conditions: processed tracks in bigWig format were downloaded from GEO databases: H3K4me1 and H3K27me3 from GSE72164^57^; H3K27ac from GSE92407^58^; ATAC-seq and H3K4me1 from GSE155062^59^. All bigWig tracks were converted to BED and lifted to the mm39 assembly using liftOver^60^ tool using corresponding UCSC chain files(mm9ToMm39.over.chain.gz, mm10ToMm39.over.chain.gz).The liftover bed files were sorted with bedtools, and converted back to bigWig using bedGraphToBigWig^61^. We normalized each track to CPM. Biological replicates for each histone mark were averaged at each position.

### Processing of STARR-Seq data

We used preprocessed STARR-Seq data generated in mESCs (GSE143544^30^). STARR-Seq signals in the form of Empirical Bayes Shrunken Enrichment (EBSE) values were lifted over to mm39 following the same procedure as was described above.

### Processing of TF ChIP-seq data

Coverage bigwig files for transcription factor ChIP-seq datasets were sourced from the ReMap database^31^. Data were processed as described in ReMap until peak-calling. Bigwig files were generated using MACS2^47^. Fragment length was determined independently using MACS2 predicted with options “--gsize mm -m 5 50”. Peaks were called using MACS2 callpeak with options “--gzize mm --SPMR –bdg –extsize $fragment_length --buffer-size 100000”. The resulting bedgraph were converted to bedgraph using UCSC bedGraphToBigWig. The pipeline is available on GitHub (https://github.com/remap-cisreg/remap-pipeline), and the processed data are deposited in the Recherche Data Gouv repository (doi.org/10.57745/GEIHVN). We used only data sets annotated as untreated or Ctrl naive mESCs: sets associated with specific genetic perturbations, targeted knockdowns, or non-zero treatments were excluded. bigWig files aligned to the mm10 assembly were converted to the mm39 assembly using the CrossMap^62^. ChIP-seq bigWig files targeting the same TF were pooled together, summing their read coverage at base pair resolution. To ensure comparability between different tracks, individual processed files were normalized by a scaling factor: base-paired resolution signal was divided by the total coverage of each respective bigwig file and multiplied by 10^8^.

### Visualization of average genomics track signal at enhancers

To evaluate the spatial distribution of genomic signals from transcription factor and histone modification ChIP-seq, ATAC seq and other genomic tracks at enhancer loci (Figs. 5G and 6A), continuous signals were extracted from the normalized bigWig files. For each DHS, signal coverage was retrieved at single-base resolution across a ±1 kilobase genomic window anchored on the DHS midpoint. For each genomic track (DNase I-Seq, ATAC-Seq, ChIPSeq of histone modifications or TF), the positional signals were averaged across all DHSs within each TSS-spacing group as defined above. For visualization this signal was normalized between 0 (observed minimum) and 1 (observed maximum) within the enhancer regions visualized.

### Sequence logo analysis around enhancer TSSs

To evaluate the sequence composition at enhancer TSSs, we analyzed the genomic sequences flanking the forward and reverse strand TSSs. For each annotated DHS, we defined a -100bp (upstream) to +20bp (downstream) genomic window anchored on both eRNA TSSs, and we extracted the corresponding DNA sequences from the reference genome in a strand-aware manner. Sequences containing non-canonical nucleotides were discarded. These sequences were then grouped according to their previously defined TSS-spacing groups. Nucleotide fractions per base position were calculated adding a pseudocount of 0.01 to each nucleotide, and visualized as sequence logos^63^ using ggseqlogo^64^.

### Analysis of activation of poised mESC enhancers in anterior neural progenitor cells

To evaluate changes of chromatin modifications and P300 occupancy between cellular states, raw ChIP-seq data for naive mESCs and differentiated anterior neural progenitor cells (AntNPCs) were downloaded from GEO: GSE89209^33^. Reads were aligned to the mm39 reference genome using Bowtie2^65^ using paired ends parameters. Filtered alignments were converted into BigWig signal tracks utilizing bamCoverage from deepTools^66^. Signal coverage was generated as bigWig files at 10-base-pair resolution extending the paired-end reads to account for the full sequenced fragment lengths. Tracks were normalized by their total coverage, as described above. To calculate changes between states, for each ChIP-seq data set enhancer, we calculated AntNPC vs mESC log_2_FC values, using the sum of reads in a ±250 base pair (500 bp wide) window relative to enhancer midpoints. For visualisation, we ranked enhancers by their *d* values and estimated the mean log_2_fold-change and the 95% confidence intervals across the ranked enhancers.

### 4sU-seq and GRO-seq analysis

BigWig coverage tracks of 4sU-seq and GRO-seq ESC-to-NPC differentiation datasets (downloaded from GEO:GSE115774^35^ (NPCs harvested at 3 days) and GEO: GSE142312^36^ (NPCs harvested at 6 days) respectively) were lifted over to the mouse mm39 reference genome using CrossMap^62^. For each data type, technical and biological replicates for each biological state (*i.e.*, 4sU-seq mESCs and NPCs(day 3); GRO-seq mESC and NPC(day 6) were pooled into combined coverage tracks. Enhancer loci were selected as described in the “Enhancer curation” section. Unlike TLDR-seq, 4sU-seq and GRO-seq measure nascent RNA fragments rather than capped transcript 5′ ends. Consequently, signals at intragenic enhancers may derive from transcription of the overlapping host gene, including transcription initiated from alternative gene promoters, rather than from the enhancer itself. To minimize this ambiguity, the analysis was restricted to intergenic enhancers. We excluded any enhancer overlapping a GENCODE-annotated protein-coding or lncRNA gene body, or a promoter-proximal region defined as the 2 kb immediately upstream of an annotated TSS. Additionally, to avoid confounding RNA originating outside the locus (i.e., Pol II read-through into the enhancer) we discarded loci whose GRO-seq or 4sU-seq read depth at the upstream enhancer locus boundaries (relative to Pol II transcriptional direction) was more than 10% of the maximum read depth from the same data type across the enhancer locus. For retained enhancer regions, transcribed blocks were defined using a threshold depending on each enhancer’s maximum signal: a main TSS was assigned to the 5’ edge of the most upstream block that retained >33% of the total 4sU-seq or GRO-seq signal coverage on the enhancer locus. We analyzed the 4sU-seq and GRO-seq datasets separately. For each, we restricted the analysis to enhancer loci exhibiting bidirectional signal (which can be either convergent or divergent) at both the starting and ending time points and we calculated the *d* metric as above for each of these loci. We then identified enhancers switching from a convergent (*d* < 0) to a divergent (*d* > 0) state during differentiation.

### Statistics and visualizations

All statistics were done in R (https://www.R-project.org/) and the tidyverse package suite^67^. Graph visualizations were made using ggplot2^68^, ggseqlogo^64^ and pheatmap (https://github.com/raivokolde/pheatmap) packages. Genome browser plots were based on the Integrative Genomics Viewer (IGV) browser^69^.

## Author contributions

YD, ASt, MP and ASa analyzed the data. JYA and CV prepared TLDR-seq data. WG and KW performed depletions, cell work and RNA extractions. LS and BB contributed newly processed ReMap data. YD, ASt, ASa, MP, AG, THJ and RA interpreted data. ASa wrote the first draft. ASa, YD, ASt,THJ, KW, AA, WG, and JYA edited the manuscript.

---

**Figure S1. Extends Figure 1:**
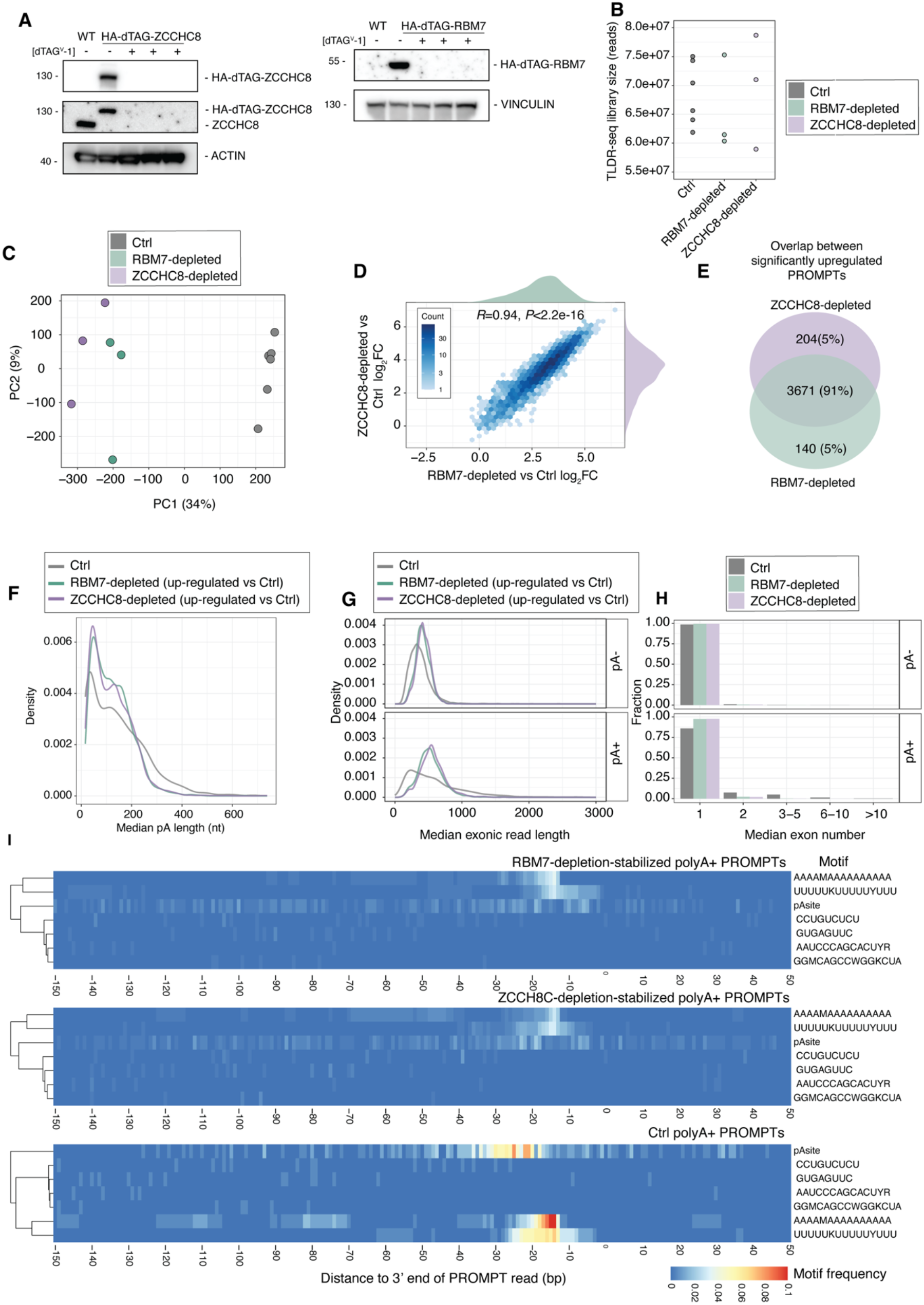
**A: Validations of ZCCHC8 and RBM7 depletions.** Western blot analysis of lysates from Ctrl (WT) and HA-dTAG-ZCCHC8 (left) and HA-dTAG-RBM7 (right) in mES cells. Samples were either treated with DMSO (-) or dTAG^V^-1 (+) for 4h prior to harvest. Membranes were probed with antibodies against HA to confirm tagged ZCCHC8 and RBM7 expression, along with antibodies against ZCCHC8 to compare with Ctrl levels. ACTIN and VINCULIN were used as loading controls. **B: Number of TLDR-seq reads in initial libraries.** The Y axis shows the number of reads. Each dot is one library. The X axis and dot color shows the experimental group. **C: Principal component (PC) analysis-based expression patterns per library on TLDR-seq cluster level.** Axes show PC1 and 2, with % variance covered indicated. Dots indicate libraries, colored by experimental group. **D: RBM7 and ZCCHC8 depletion sensitivity of TLDR-seq 5’ clusters.** X and Y axes show log_2_FC values of ZCCHC8 and RBM7 depletions vs Ctrl, respectively, for each TLDR-seq 5’ cluster. Distributions of each set of log_2_FC values are shown on top or right of the plot. Hexagons are colored by the number of observations as indicated in the legend. Pearson’s R and correlation test *P* value are shown. **E: Overlap in TLDR-seq PROMPT clusters significantly upregulated by RBM7- or ZCCHC8-depletion.** Venn diagram shows overlap of upregulated PROMPT TLDR-seq clusters vs Ctrl: number of clusters in each Venn diagram region, and percentage of union are shown. **F: Distribution of median pA length of reads per PROMPT TLDR-seq cluster.** The Y axis shows probability density. The X axis shows the median pA length of pA^+^ reads within a PROMPT TLDR-seq cluster. Colors as in panel 1D. Only upregulated clusters (log_2_FC >1 and *FDR*<0.05 vs. Ctrl) are shown for RBM7 and ZCCHC8-depletion data, while Ctrl data shows all clusters. **G: Distribution of median exonic length of reads per PROMPT TLDR-seq cluster.** The Y axis shows probability density. The X axis shows the median total exonic length of TLDR-seq reads within a PROMPT TLDR-seq cluster. pA^+^ and pA^-^ reads are analyzed separately. Colors as in panel 1D. Only upregulated clusters (log_2_FC >1 and *FDR*<0.05 vs Ctrl) are shown for RBM7 and ZCCHC8-depletion data, while Ctrl data shows all clusters. **H: Distributions of median number of exons per PROMPT TLDR-seq cluster.** The Y axis shows the fraction of clusters. The X axis shows the median number of exons from reads assigned to a TLDR-seq cluster. Colors as in panel 1D. Data is separated by polyA+ and polyA- reads. Only upregulated clusters (log_2_FC >1 and *FDR*<0.05 vs Ctrl) are shown for RBM7 and ZCCHC8-depletion data, while Ctrl data shows all clusters. **I: Sequence motif occurrence around polyadenylated PROMPT 3’ ends reads.** X axis as in Fig. 1H. Each row corresponds to one sequence motif as shown on the right, identified by pattern finding algorithms where color indicates motif occurrence probability at respective position. Top heat maps show polyA+ PROMPT reads that were stabilized upon depletions. The bottom heat map shows Ctrl pA^+^ PROMPT reads.

**Figure S2: extends Figure 4.**
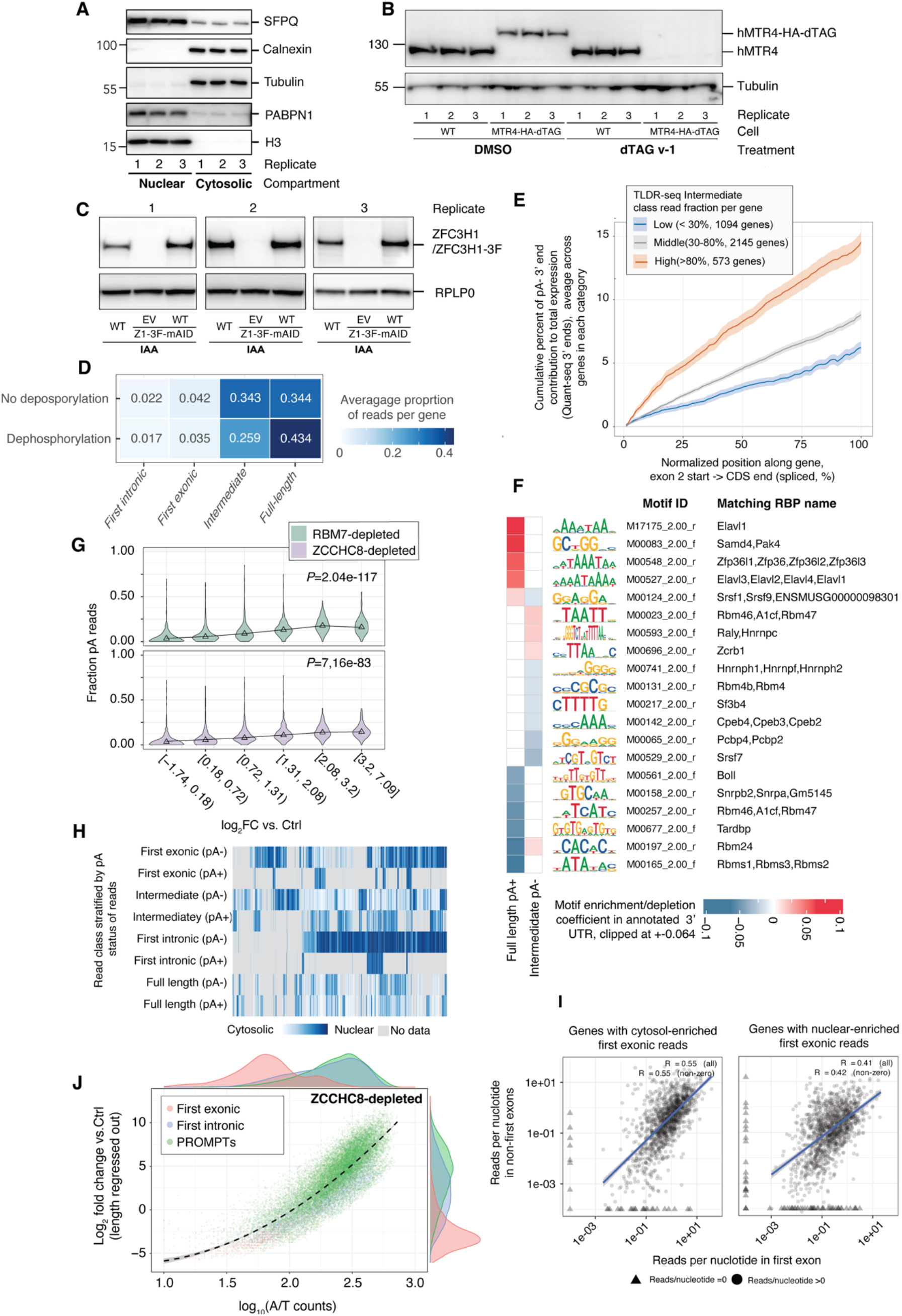
**A: Cytosolic/nuclear extraction validation.** Western blot confirming enrichment of SFPQ, PABPN1 and histone H3 in the nuclear fraction, and Tubulin and Calnexin (ER) in the cytoplasmic fraction. **B: hMTR4 depletion validation.** Western blot confirming depletion of hMTR4-HA-dTAG following 4h treatment with dTAG^V^-1. **C: ZFC3H1 deletion validation.** Western blot confirming expression of ZFC3H1-3F in the wild type overexpression cell line but not the empty vector cell line following rapid depletion of endogenously tagged ZFC3H1-3F-mAID. **D: Average fraction of genic TLDR-seq 3’ ends falling into specific classes as a function of TLDR-seq dephosphorylation step inclusion.** Rows show results from TLDR-seq on wild-type mES cells with or without the 3’ dephosphorylation step included. For each gene (1805 multi-exonic genes having >0 reads in each sample), we calculated the fraction of reads that fell into a certain read class (column) and then calculated the average fraction across all genes. This average number is shown in each box. Box color also indicates the average fraction. Data is based on three replicates in each row. **E: QuantSeq 3’ end analysis confirms that degradation intermediates scale with the TLDR-seq intermediate fraction.** The Y axis shows cumulative contribution in percent of QuantSeq pA− 3′ ends along the spliced region from the start of exon 2 to the CDS end. The X axis shows the normalized position of 3’ reads within the spliced RefSeq model, where 0 is the start of exon 2 and 100 is the end of the CDS, using 100 equally spaced bins. For pA- vs pA+ distinction and scaling, see Methods. Genes are grouped by their TLDR-seq intermediate pA− fraction: < 30% (blue, *n* = 991), 30–80% (grey, *n* = 1,909) and > 80% (orange, n = 511). Lines show the mean per group and ribbons show ± s.e.m. **F: 3’ UTR RNA binding protein motif enrichments for Full-length and Intermediate classes.** Regression coefficients plot visualizing top 3’UTR RBP binding predictors for proportion of full-length pA^+^ RNA and intermediate pA^-^ RNA in Ctrl cells. We used the Centered Log-Ratio (CLR) of each RNA class as an enrichment/depletion indicator, which was regressed by the density of RBP binding motifs, indicated by heat map cell color. Red: RNA proportion tends to increase when the density of an RBP binding motif increases in 3’ UTR. Blue: RNA proportion tends to decrease when the density of an RBP binding motif increases in 3’ UTR. RBP motifs from CISBP-RNA^28^. Sequence logos of motifs are shown. **G: Relation between polyadenylation and stabilization following RBM7/ZCCHC8 in the First-intronic 3’ class.** The plot is organized as Fig. 2B (top) but shows analysis of reads from the First-intronic read class. **H: Heat map visualization of nuclear-cytoplasmic ranks of genic reads.** This is an extended version of Fig. 4F, following the same conversions but including all genes and read classes, where reads are separated by pA status. Grey color indicates cells with insufficient TLDR-seq reads to calculate cytosolic/nuclear ratios. **I: Relation between reads/nucleotide expression for reads ending in first exon vs. remaining exons.** The X axis shows the number of Ctrl TLDR-seq reads ending in the first exon, per exonic nucleotide, in log scale. The Y axis shows the number of Ctrl TLDR-seq reads ending in the remaining, per exonic nucleotide, in log_10_ scale. Analyses are divided depending on overall the nuclear-cytosolic bias of reads ending in the first exon, as in Fig. 4F,G. Dots indicate each first vs. remaining exon pair for each gene in each group. Triangle-shaped dots indicate that the X and/or Y axis has no reads (=0 reads), clamped at 3.5 e-05. Pearson’s *R* for each plot is shown, including 0 count observations in calculations or not. Blue lines indicate trend lines. **J: Linear model of ZCCHC8-depletion sensitivity for First-exonic, First-intronic and PROMPT reads.** The plot is organized as Fig. 4H, but uses ZCCHC8 depletion data.

**Figure S3: extends Figure 5.**
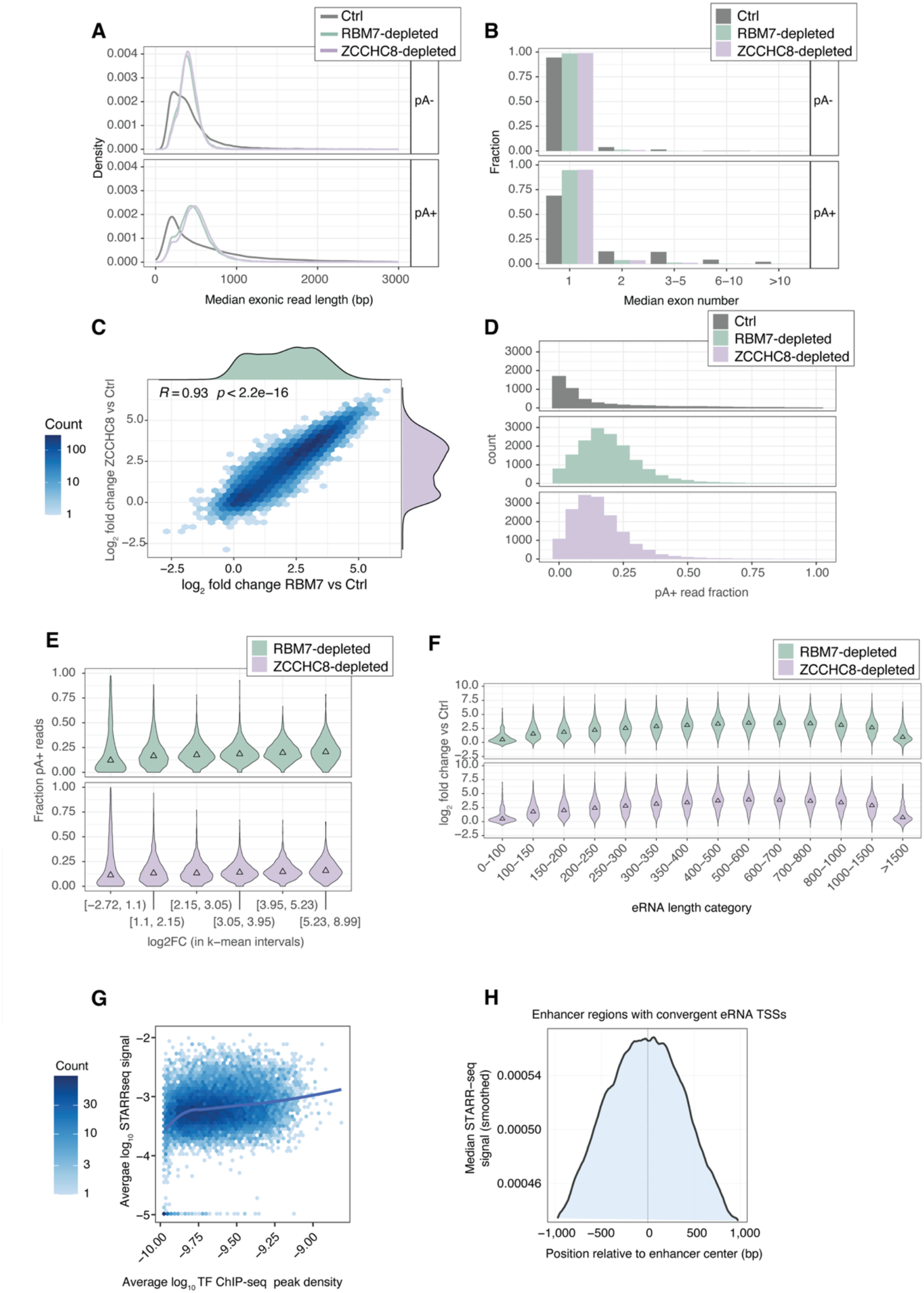
**A: Distribution of median exonic length of reads per eRNA TLDR-seq cluster.** Organized as in Fig. S1G but analyzing eRNA TLDR-seq data. **B: Distributions of median number of exons per eRNA TLDR-seq cluster.** Organized as in Fig. S1H but analyzing eRNA TLDR-seq data. **C: RBM7 and ZCCHC8 depletion sensitivity of eRNA TLDR-seq 5’ clusters.** Organized as in Fig. S1D but analyzing eRNA TLDR-seq data. **D: Distributions of fraction pA^+^ TLDR-seq reads from eRNA clusters.** Organized as in Fig. 1D but analyzing eRNA TLDR-seq data. **E: Relation between eRNA polyadenylation and stabilization following RBM7/ZCCHC8 depletion.** Organized as in Fig. 1E but analyzing eRNA TLDR-seq data. **F: Relation between eRNA length and sensitivity to RBM7 and ZCCHC8 depletions.** Organized as in Fig. 2B (top), but analyzing eRNA TLDR-seq data **G: Relation between TF ChIP-seq peak density and STARR-seq signal at enhancers.** The plot is organized as in Fig.5C but the Y axis shows log_10_-scaled ChIP seq binding peaks per enhancer nucleotide based on ReMap data. **H: Distribution of STARR-seq signal across convergent enhancer loci.** The Y axis shows median STARR-seq signal across all enhancers where *d*<0. X axis shows distance to enhancer midpoint in bp. The line is smoothed.

**Figure S4, extends Figure 6.**
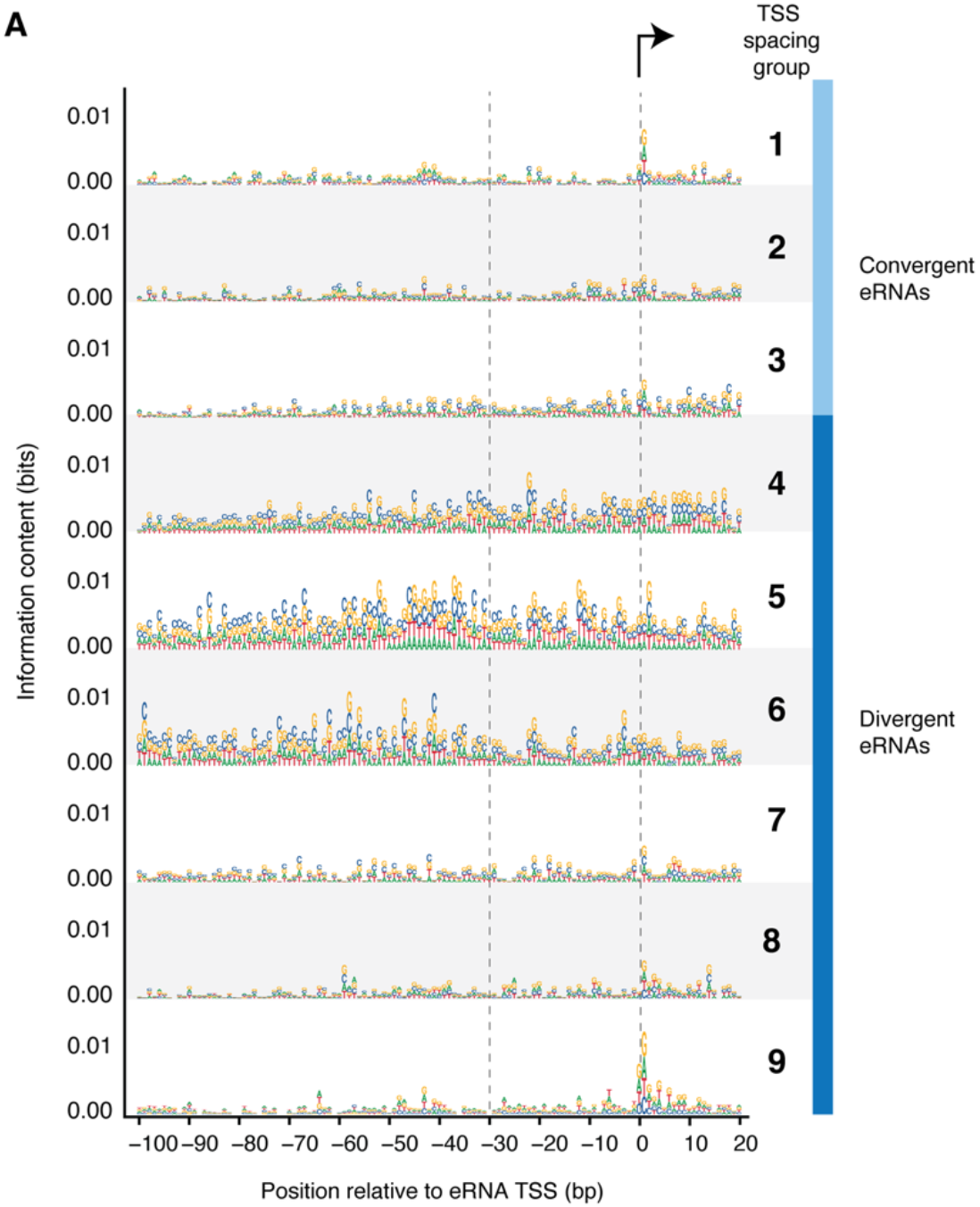
**A: Sequence patterns around eRNA TSSs stratified by eRNA spacing and orientation.** Sequence logos for each TSS spacing group, as defined in Fig. 5E, are shown. The X axis shows relative distance to positions around eRNA TSS (X=0). Transcription goes left to right. Y axis shows information content in bits: note that the Y scale is capped at 0.1 (of max 2 bits).

